# Evolution of olfactory receptors tuned to mustard oils in herbivorous Drosophilidae

**DOI:** 10.1101/2019.12.27.889774

**Authors:** Teruyuki Matsunaga, Carolina E. Reisenman, Benjamin Goldman-Huertas, Philipp Brand, Kevin Miao, Hiromu C. Suzuki, Kirsten I. Verster, Santiago R. Ramírez, Noah K. Whiteman

## Abstract

The diversity of herbivorous insects is attributed to their propensity to specialize on toxic plants. In an evolutionary twist, toxins betray the identity of their bearers when herbivores co-opt them as cues for host-plant finding, but the mechanisms underlying this process are poorly understood. We focused on *Scaptomyza flava*, an herbivorous drosophilid specialized on isothiocyanate (ITC)-producing (Brassicaceae) plants, and identified *Or67b* paralogs that were triplicated as mustard-specific herbivory evolved. Using heterologous systems for the expression of olfactory receptors, we found that *S. flava* Or67bs, but not homologs from microbe-feeding relatives, responded selectively to ITCs, each paralog detecting different ITC subsets. Consistent with this, *S. flava* was attracted to ITCs, as was *Drosophila melanogaster* expressing *S. flava* Or67b3 in the homologous *Or67b* olfactory circuit. Thus, our results show that plant toxins were likely co-opted as olfactory attractants through gene duplication and functional specialization (neofunctionalization and subfunctionalization) in drosophilid flies.

## INTRODUCTION

Many plant molecules used in food, agriculture and medicine first evolved as defenses against natural enemies ^1^. Among the most familiar are reactive electrophiles that produce pain when eaten, including diallyl disulfide and thiosulfinates in Alliaceae (*e.g.*, garlic), α, β-unsaturated aldehydes in Lauraceae (*e.g.*, cinnamon), and isothiocyanates (ITCs) in Brassicaceae (*e.g.*, arugula, radish, and wasabi). These electrophiles activate the ‘wasabi taste receptor’ TrpA1, which is conserved in flies and humans ^2^. Although ITCs are potent natural insecticides aversive to most insects, including *D. melanogaster* ^3^, some insect species are specialized on ITC-bearing Brassicaceae (mustards). This insect-plant interaction has been important in advancing the field of co-evolution ^4^. Brassicaceae specialists from many insect orders (*e.g.*, Diptera ^5^, Heteroptera ^6^, Hemiptera ^7^, and Lepidoptera ^8^) can be trapped with ITC baits in crop fields, revealing an evolutionary twist of fate in which mustard specialist insects use ancestrally aversive electrophiles as olfactory cues for host-plant finding ^9^. However, the evolutionary mechanisms underlying the chemosensory adaptations of insect herbivores to defensive plant compounds are widely unknown ^10^. Identifying these mechanisms can help us understand the evolution of herbivorous insect species, 90% of which are specialized on a limited set of host plants ^11^.

*Scaptomyza* represents a compelling genus to investigate how specialist herbivores evolved to co-opt plant defenses as host finding cues. Phylogenetically nested within *Drosophila*, *Scaptomyza* is the sister group to the Hawaiian *Drosophila* radiation ^12^, and contains many herbivorous species with varying degrees of specialization on Brassicaceae and Caryophyllaceae ^13^. Adult females make feeding punctures in living leaves using sclerotized and dentate ovipositors ^14^, and their larvae hatch directly into the mesophyll tissue, which they mine ^15^, an unusual life history within the Drosophilidae. *S. flava* specializes on Brassicaceae ^16^ and, like humans and *D. melanogaster*, uses the mercapturic pathway to detoxify ITCs ^17^. *S. flava* was first reported from *Arabidopsis thaliana* in North America as *S. flaveola* ^18^. Thus, genomic and genetic tools of both *Arabidopsis* and *Drosophila* can be utilized to dissect both sides of the plant-herbivore equation. Herbivory evolved ca. 10-15 million years ago in *Scaptomyza* (Figure 1A) and so it provides an unusually useful context to understand the evolutionary and functional mechanisms underlying chemosensory specialization, because the major herbivorous insect radiations (*e.g.* Lepidoptera^19^, Phytophaga^20^) are much more ancient in origin.

**Figure 1.**
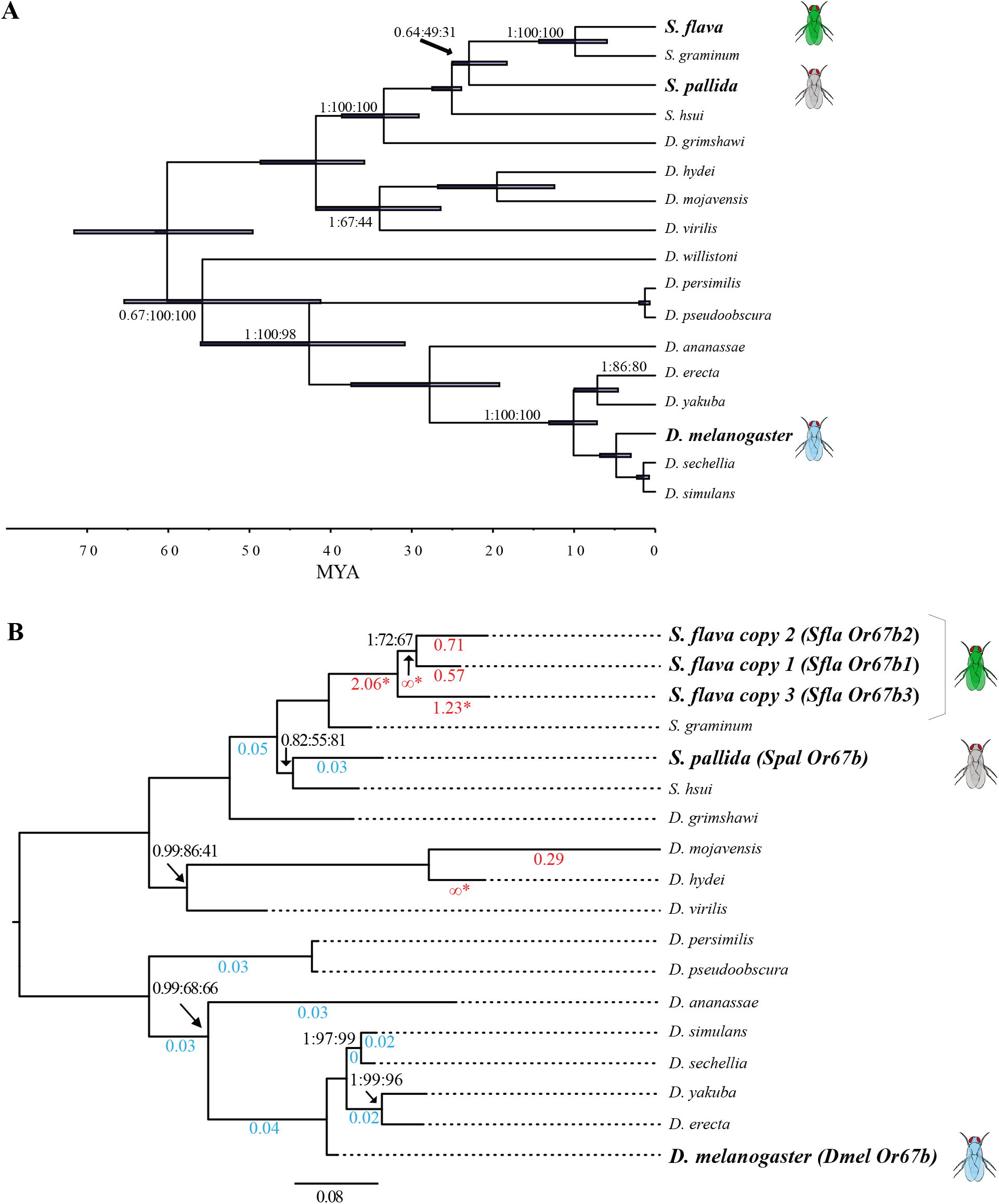
Maximum likelihood (ML) phylogeny of *Or67b* in Drosophilidae. **(A)** Time-calibrated Bayesian chronogram of *Scaptomyza* and *Drosophila* spp. inferred from nine protein coding and two ribosomal genes. Source species for the *Or67b* coding sequences crossed into the empty neuron system are indicated with fly pictograms. Bars indicate 95% highest posterior density (HPD) age estimates in millions of years ago. Tree topology labeled as follows: Posterior Probability (PP):ML BS:Parsimony BS on branches. Scale bar proportional to MYA. (**B)** ML phylogeny reconstructed based on the coding sequence of *Or67b* orthologs from twelve *Drosophila* species, *S. pallida, S. hsui, S. graminum,* and *S. flava*. All bootstrap supports for the nodes are >80% and all posterior probabilities were >0.95 for the MrBayes tree. Branches with significant support (FDR p-value < 0.05) for d*_N_*/d*_S_* values different from the background rate are indicated with colored branch labels (blue where the foreground rate is less than the background, and red/enlarged fonts where d*_N_*/d*_S_* is greater than the background). Only *S. flava* and *D. mojavensis* branches have significantly elevated d*_N_*/d*_S_* according to branch model tests. *S. flava*, *S. pallida* and *D. melanogaster* labeled identically to (A). Scale bar units are substitutions per site.

*S. flava* lost three olfactory receptor (*Or*) genes encoding canonical yeast-associated receptors after divergence from microbe-feeding drosophilid relatives, contemporaneous with the evolution of herbivory ^21^. However, the loss of function does not explain how herbivorous *Scaptomyza* species evolved attraction to Brassicaceae plants (a gain of function phenotype).

Gain of function through gene duplication and subsequent divergence plays an important role in evolutionary innovation and is often associated with trophic transitions ^22^. Gene and whole genome duplications in Brassicaceae in the last 90 million years resulted in the evolution of glucosinolates, the precursors of ITC compounds ^23^. Reciprocally, in diverse Brassicaceae-specialist pierid butterflies, enzymes that divert hydrolysis of aliphatic glucosinolates away from ITC production to less toxic nitriles evolved in tandem, underpinning their diversification and highlighting the importance of gene family evolution in this co-evolutionary arms race ^24^. While our understanding of detoxification in plant-herbivore systems has grown rapidly over the past decade ^25^, the evolutionary mechanisms underlying the chemosensory basis of hostplant orientation and finding remain relatively unknown. Here we investigated the extent to which duplication of chemosensory genes in the *S. flava* lineage contributed to attraction to specific host-plant volatiles, including ITCs.

We took a genes-first approach to study olfactory host-plant specialization in *S. flava*. Three chemoreceptor protein families (olfactory receptors -Ors-, ionotropic receptors -Irs-, and nociceptive receptors -Trps-) are candidates for mediating responses to host-specific volatile electrophiles such as ITCs ^26^. In particular, insect Ors are collectively sensitive to a variety of odorants important for food and host finding, avoidance of predators, and animal communication, including aversive chemicals ^27^, pheromones ^28–30^, and host-, oviposition-, and food-related attractive compounds ^31–33^. Accordingly, we scanned the genome sequence of *S. flava* to identify rapidly evolving Ors and found a lineage-specific gene copy number expansion of the olfactory receptor gene *Or67b* that we named *Or67b1, Or67b2,* and *Or67b3*. The coding sequences of these three paralogs exhibit signatures of positive natural selection and duplicated likely within the last ca. 15 million years, at the base of the mustard-feeding clade. In contrast, *Or67b* is present as a single copy under strong evolutionary constraint across the Drosophilidae ^21, 34–35^, including two focal species included here, the microbe-feeding close relative *S. pallida* (subgenus *Parascaptomyza*), and the more distantly related *D. melanogaster*. Our *in vivo* functional characterizations of all full-length Or67b proteins from these three species show that the three *S. flava* Or67b proteins, but not the conserved Or67b proteins from its microbe-feeding relatives, responded selectively and sensitively to volatile ITCs. Concomitantly, we found a population of *S. flava* antennal olfactory sensory neurons (OSNs) responsive to ITCs. In agreement with these results, *S. flava*, but not *S. pallida* or *D. melanogaster*, is attracted to odors from Brassicaceae plants, including single ITC compounds. Expression of *Sfla* Or67b3 in the *D. melanogaster* homologous Or67b olfactory circuit indicates that *Sfla* Or67b3 can confer odor-oriented responses towards ITCs. Finally, our genetic silencing experiments demonstrate that the Or67b olfactory circuit mediates attraction in *D. melanogaster*. These findings support the hypothesis that simple evolutionary changes in the odorant tuning of an Or may be sufficient for changing the identity of the odorants that evoke behavioral responses ^28, 36–38^. Of particular ecological significance is that the three *Sfla* Or67b proteins shifted the odor-receptive ranges from the typical ancestral Or67 drosophilid response profile of ketones, alcohols, and aldehydes ^39–40^ to ITCs, an entirely different chemical class of compounds. In summary, our results suggest a relatively simple sensory mechanism by which mustard-specialist herbivorous insects may have evolved olfactory attraction towards host-plant ITCs, via gene duplication followed by neofunctionalization (see discussion) of a specific clade of otherwise highly conserved Ors in the Drosophilidae.

## RESULTS

### Phylogenetic analysis identified *Or67b* paralogs as candidates mediating the detection of ecologically relevant host-plant odorants

To search for candidate ITC-detecting chemosensory receptors in *S. flava*, we conducted a phylogenetic analysis of the Or protein sequences of *S. flava* and four other *Drosophila* species to identify *Scaptomyza*-specific gene losses and gains (Figure supplement 1) as well as those that were rapidly evolving at the protein level, regardless of duplication history. The Or topology was largely congruent with previous *Drosophila* Or gene trees ^21^, except for some deeper nodes, which had low bootstrap support. In addition to a previous analysis of *Or* coding sequences using a branch-site test to identify *Scaptomyza Or* codons likely evolving under positive selection ^21^, we also fit simpler foreground/background branch models, as implemented in PAML, to scan for *Ors* whose sequences were evolving under divergent selection regimes from the *Drosophila* background.^41^ Out of seventy-five *S. flava* branches tested, seven branch models, corresponding to paralogs of *Or63a, Or67b* and *Or98a* in *D. melanogaster*, inferred a foreground rate larger than one, consistent with a high rate of fixation of nonsynonymous mutation during positive selection across the *Or* coding sequence (supplementary file 1). Among these receptors, *Or63a* is only expressed in *D. melanogaster* larvae ^42^, and the *Or98a-*like genes found in *Scaptomyza* have no *D. melanogaster* homologs and have not been characterized functionally. In contrast, *Or67b* modulates oviposition behavior in adult *D. melanogaster* ^43^, and is implicated in selection of oviposition sites in *D. mojavensis* ^44^, making it a good candidate for the olfactory adaptation of *Scaptomyza* to a novel host niche. After expanding the representation of *Or67b* orthologs in a phylogenetic analysis and conducting branch tests in PAML on all single branches on the *Or67b* phylogeny, we found significantly elevated d*_N_*/d*_S_* exclusively in *S. flava* and *D. mojavensis Or67b* (Figure 1B and Supplementary file 1). Furthermore, synteny analysis confirmed that the regions upstream and downstream of the two non-syntenic *Or67b* copies in *S. flava* are present in both *S. pallida* and *D. melanogaster*. Thus, the absence of each paralog in *S. pallida* and *D. melanogaster* is not a genome assembly artefact but rather, an actual absence (Figure supplement 2A, B).

### *S. flava* Or67bs respond specifically to mustard plant odors

Because *Or67b* is triplicated exclusively in the *S. flava* lineage and likely evolved under positive selection since its divergence from copies in non-herbivorous species, we next investigated whether *Sfla* Or67bs acquired novel ligand-binding sensitivity towards odorants characteristic for the mustard hosts of *S. flava* (Figure 1). First, we confirmed that all three *S. flava* paralogs and the *S. pallida* ortholog are expressed in adults (Source data). To study the odor-response profile of Or67b across species, we then heterologously expressed the Or67b paralogs in the *D. melanogaster* olfactory at1 or ab3A neurons lacking endogenous Ors ^28, 45^, and measured electrophysiological responses to an array of mustard secondary plant compounds and other odors (Figure 2A and Figure supplement 3A). Ab3A neurons expressing *Sfla* Or67b1*, Sfla* Or67b3*, Spal* Or67b, and *Dmel* Or67b showed spontaneous bursts of action potentials (as described for other Ors expressed in these neurons ^46^), whereas neurons expressing *Sfla* Or67b2 showed spontaneous activity only when expressed in at1 OSNs. Thus, we used the “at1 empty neuron” system for studying the olfactory responses of *Sfla* Or67b2, and the “ab3A empty neuron” system for investigating the responses of all other Or67b proteins.

**Figure 2.**
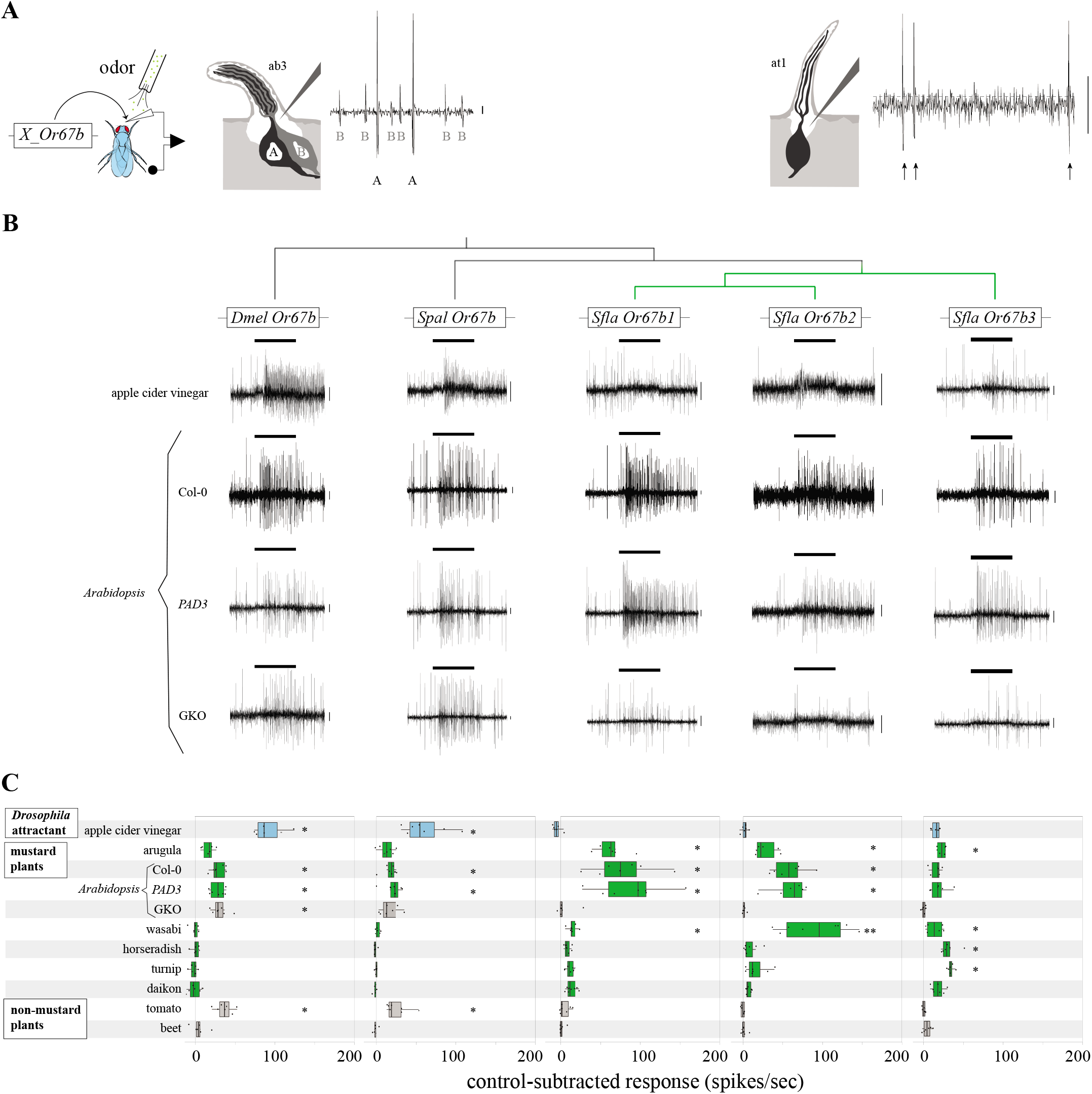
Responses of homologs Or67bs from *D. melanogaster*, *S. pallida*, and *S. flava* expressed in the *D. melanogaster* empty neuron systems to stimulation with natural odor blends. **(A)** Schematic representation of the single sensillum recording (SSR) using two “empty neuron systems”. Or67b proteins (*X_Or67b*, where *X* refers to the fly species) were expressed in a *D. melanogaster* mutant that lacks its endogenous *Or22a* in ab3A (antennal basiconic 3A) OSNs ^97^, or *Or67d* in at1 (antennal trichoid 1) OSNs ^28^. Note that the at1 empty neuron system was used only for expression of *Sfla* Or67b2, as this Or was not functional in the ab3A empty neuron system. In the antennal basiconic empty neuron system (left) the sensilla houses the A neuron (which expresses one of the Or67b proteins) and the native intact B neuron. The A neuron has larger amplitude spikes than the B neuron, allowing separation of spikes originating from either of them. The antennal trichoid empty neuron system houses a single OSN expressing *Sfla* Or67b2 (see also Figure supplement 3). Calibration bars: 10 mV throughout all figures unless otherwise noted. **(B)** Representative electrophysiological SSRs obtained from the targeted sensilla of flies expressing Or67b in OSNs in response to stimulation with apple cider vinegar, wild-type Col-0 *Arabidopsis thaliana*, and *PAD3* and quadruple aliphatic and indolic glucosinolate knockout (GKO; *CYP79B2, CYP79B3, MYB28, MYB29*) *A. thaliana* mutant lines. Although all three *A. thaliana* genotypes have the same genetic background ^50–51^, *PAD3* plants are a more appropriate control for GKO than Col-0, since *PAD3* is deficient (as is GKO) in the production of camalexin but not aliphatic or indolic glucosinolates. The bars above records indicate the onset and duration (1 sec) of the stimulation throughout all figures unless otherwise noted. **(C)** Responses (net number of spikes/second, control-subtracted, n=6-9 obtained from 2-4 animals) evoked by stimulation with apple cider vinegar, mustard leaf odors (arugula and *A. thaliana*), mustard root odors (wasabi, horseradish, turnip and daikon), non-mustard leaf odors (tomato), and non-mustard root odors (beet, control). The outer edges of the horizontal bars represent the 25% and 75% quartiles, the vertical line inside the bars represents the median, and the whiskers represent 10% and 90% quartiles; each dot represents an individual response. Asterisks indicate significant differences between the control-subtracted net number of spikes and a threshold median value (10 spikes/second), as explained in material and methods (one-sample signed rank tests; * p<0.05, **p<0.01). Neurons expressing *Dmel* Or67b and *Spal* Or67b, but not those expressing any of the *S. flava* Or67b paralogs, responded to stimulation with apple cider vinegar. Conversely, only neurons expressing *S. flava* Or67b paralogs responded to arugula odors (which bear ITCs). *Dmel* Or67b responded to all *A. thaliana* genotypes, while *Sfla* Or67b1-2 responded only to ITC-bearing *A. thaliana*, indicating that the presence of ITCs within plants is necessary to evoke responses from these two *S. flava* paralogs. Stimulation with wasabi root odors evoked responses from all *Sfla* Or67b paralogs but not from the *Dmel* or the *Spal* paralogs.

In order to test which odors specifically activate *Sfla* Or67b paralogs, we tested the responses of all Or67bs to stimulation with apple cider vinegar (a potent *D. melanogaster* attractant ^47^), crushed arugula leaves (*Eruca vesicaria*, a mustard), and crushed tomato leaves (*Solanum lycopersicum*, a non-mustard plant that releases large quantities of volatile organic compounds ^48^). OSNs expressing *Dmel* Or67b and *Spal* Or67b, but not those expressing any of the *Sfla* Or67 paralogs, responded strongly to stimulation with apple cider vinegar (Figure 2B, C), indicating that the *Sfla* Or67b paralogs lost the ancestral odorant sensitivity. Volatiles from crushed tomato leaves activated both *Dmel* Or67b and *Spal* Or67b. In contrast, all *Sfla* Or67b paralogs responded to volatiles from crushed arugula leaves, but not to those from tomato (Figure 2C), suggesting that *Sfla* Or67bs specifically respond to volatiles characteristic of their hosts but not to those from non-hosts.

In contrast to tomato plants, Brassicaceae plants, including arugula, produce glucosinolates, some of which are hydrolyzed into ITCs upon tissue damage ^49^. To test if responses of *Sfla* Or67bs to mustard leaf odors are mediated by ITCs and/or by other mustard volatile plant compounds, we used crushed leaves of wild-type *A. thaliana* (Col-0) and two loss of function mutants generated in the Col-0 background. One of these mutants (GKO) is deficient in the production of ITC-precursors derived from aliphatic and indolic glucosinolates as well as camalexin ^50–51^ and does not release ITCs upon wounding^50, 52–54^, whereas the control line (PAD3) produces these glucosinolates but not camalexin ^51, 54^. Stimulation with both wild-type *A. thaliana* and *PAD3* mutant leaves, which can produce ITCs, activated OSNs expressing *Sfla* Or67b1, b2 and b3, whereas stimulation with GKO mutant leaves did not (p>0.05 in all cases; Figure 2C). This suggests that ITCs, but not other leaf volatiles, activate the three *Sfla* paralogs. In contrast, *Dmel* Or67b expressing OSNs showed similar responses to all three *A. thaliana* genotypes (Figure 2C), in agreement with the reported responses of this Or to green leaf volatiles (GLVs) ^55^. Similar to *Dmel* Or67b, OSNs expressing *Spal* Or67 responded to all *A. thaliana* genotypes, but with low spike frequency (Figure 2C). Because GKO plants differed from both *PAD3* and Col-0 plants only in the capability of producing ITCs ^50–51^, these results demonstrate that the *Sfla* Or67b paralogs are selectively activated by these signature chemical compounds of Brassicaceae. In contrast, *Dmel* and *Spal* Or67b were activated by non-ITC mustard and non-mustard plant volatiles, consistent with previous studies ^56–57^.

Because mustard plant roots release a variety of volatile organic compounds (VOCs) including ITCs, but not GLVs ^58–60^, we also used preparations of these tissues as stimuli. We prepared root homogenates of four mustard plant species, including wasabi (*Eutrema japonicum*)^58^, horseradish (*Armoracia rusticana*)^58^, turnip (*Brassica rapa*)^59^, daikon (*Raphanus sativus*)^60^, and a non-mustard control root vegetable species (beet, *Beta vulgaris*)^61^ from a different plant order. Interestingly, OSNs expressing each *Sfla* Or67b paralog differed in responsiveness to mustard plant leaf and root volatiles. For example, stimulation with arugula elicited strongest responses in neurons expressing *Sfla* Or67b1 (64 net spikes/sec; median; Figure 2C), stimulation with wasabi root odors produced strongest responses in neurons expressing *Sfla* Or67b2 (96 net spikes/sec), and stimulation with turnip roots produced the strongest responses in OSNs expressing *Sfla* Or67b3 (34 net spikes/sec).

### *S. flava* Or67b paralogs have different ITC selectivity

We next investigated the odor-tuning profiles of Or67b copies from all tested species by testing their responses to a panel of 42 individual odorants (all tested at 1:100 vol/vol), which included ITCs, nitriles, GLVs, and odorants known to activate *Dmel* Or67b, including ketones, esters, and alcohols ^55^; Figure 3B). Testing this broad array of odors from diverse chemical classes allowed us to cover a wide range of known secondary plant compounds and other appetitive odors to investigate the odor-response profiles of Or67b proteins across species ^10, 62^.

**Figure 3.**
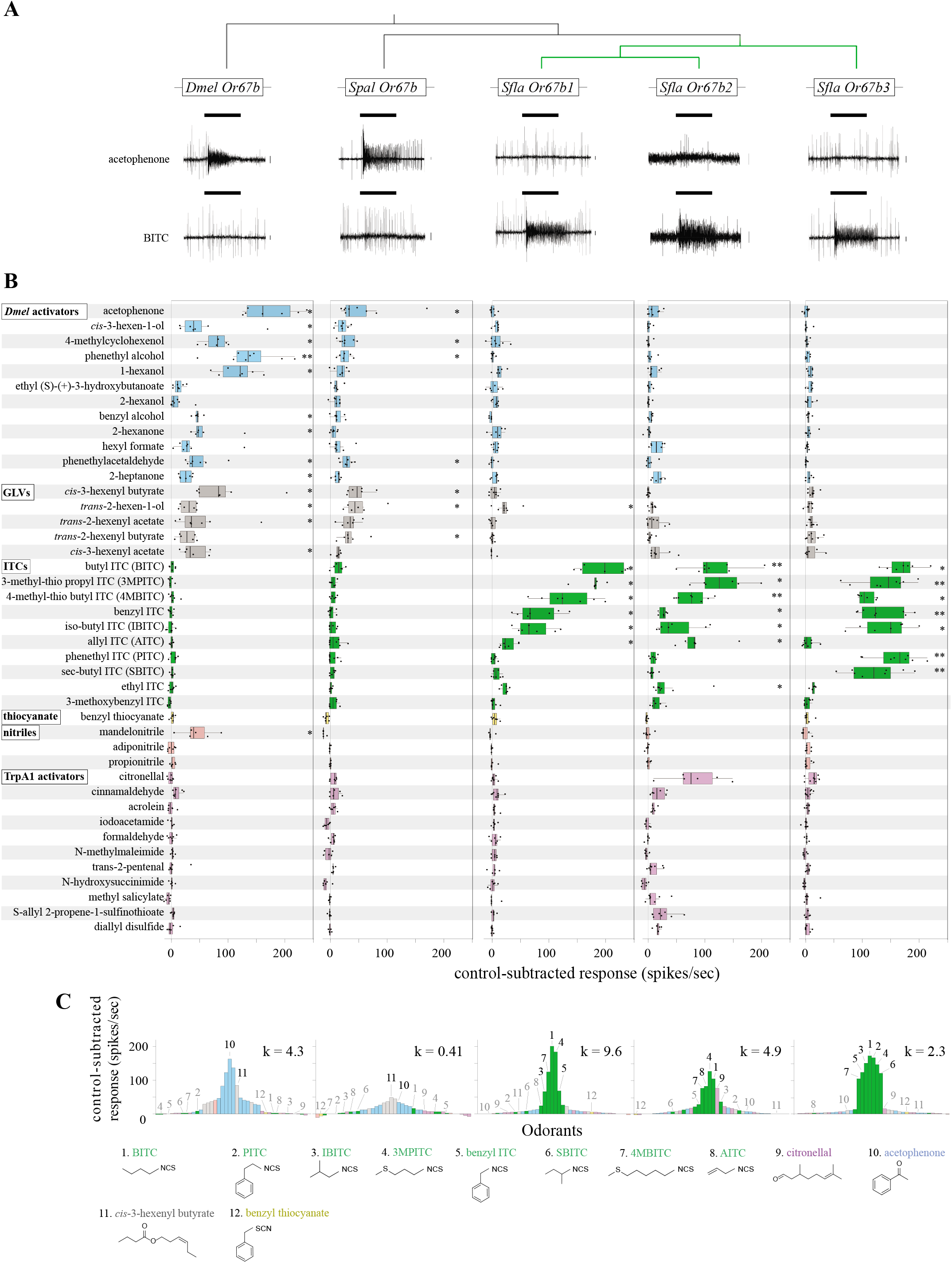
Responses of homologs Or67bs from *D. melanogaster*, *S. pallida* and *S. flava* expressed in the *D. melanogaster* empty neuron systems to stimulation with single odorants. Experiments were conducted and analyzed as in Figure 2. As before, the at1 empty neuron system was only used for expressing *Sfla* Or67b2. **(A)** Representative electrophysiological recordings obtained from the targeted sensilla of flies expressing *Or67b* genes in the empty neuron in response to stimulation with acetophenone and BITC at 1:100 vol/vol. **(B)** Responses evoked by stimulation with single odorants (tested at 1:100 vol/vol) categorized as follows: *Dmel* Or67b activators (Database of Odor Responses ^55^; blue), green leaf volatiles (GLVs; gray), ITCs (green), benzyl thiocyanate (yellow), nitrile (pink), and TrpA1 activators (purple). OSNs expressing any of the *Sfla* Or67b paralogs respond strongly and selectively to ITCs. Note that OSNs expressing any of the *S. flava* paralogs do not respond to benzyl thiocyanate stimulation (yellow bars), indicating that the presence of the ITC functional group (-N=C=S) is necessary for evoking responses from these paralogs. *Spal* Or67b and *Dmel* Or67b have similar odor-response profiles, responding mostly to stimulation with *D. melanogaster* activators and GLVs (*p<0.05, **p<0.01, one-sample signed rank tests performed as explained in the caption to Figure 2). Most odorants were diluted in mineral oil but a few needed to be prepared in other solvents (see material and methods); spikes count in response to control solvent applied in each case were subtracted from odorant-evoked responses. **(C)** Tuning curves for each Or67b, showing the distribution of median responses to the 42 odorants tested (color-coded as in **A**). The odorants are displayed along the horizontal axis according to the net responses they elicit from each Or. The odorants (numbers) eliciting the strongest responses for each Or are located at the center of the distribution and weaker activators are distributed along the edges. Note that the strongest responses (center of the distribution) from *Dmel* Or67b and *Spal* Or67b are evoked by *D. melanogaster* activators and GLVs (blue and gray bars), while the strongest responses from all *Sfla* Or67b paralogs are evoked by ITCs (green bars). The tuning breadth of each Or is quantified by the kurtosis value (k) of the distribution ^104^, with higher values indicating narrower odor-response profiles. The chemical structure of the top seven *Sfla* Or67b3 activators, as well as AITC, citronellal, acetophenone, *cis*-3-hexenyl butyrate and benzyl thiocyanate (BTC) are shown at the bottom.

We first verified the known *Dmel* Or67b response profile ^55^, which includes many chemicals from a diverse number of chemical classes: 13/17 odorants categorized as *Dmel* activators and GLVs evoked responses (one-sample signed rank tests, p<0.05; Figure 3). *Spal* Or67b responded to a smaller number of odorants within those categories (7/17 odorants; p<0.05). Strikingly, only one of the compounds that elicited responses from *DmelOr67b* and *Spal* Or67b (*trans*-2-hexen-1-ol) evoked responses from *Sfla* Or67b1 (median=22 net spikes/second; p<0.05), but none of the 17 odorants categorized as *Dmel* activators and GLVs evoked responses from *Sfla* Or67b2 or *Sfla* Or67b3 (Figure 3B; p>0.05 in all cases). While various ITCs evoked responses from each of the *S. flava* paralogs (7/10 ITCs for all three paralogs; p<0.05), none evoked responses from *Dmel* Or67b or *Spal* Or67b (p>0.05; Figure 3A, B). Tuning curves (Figure 3C) of all Or67b proteins show that *Sfla* Or67bs have response profiles distinct from those of *Dmel* Or67b and *Spal* Or67b, whereas non-ITC compounds evoke the strongest responses (acetophenone and *cis*-3-hexenyl-butyrate, respectively, center of the distribution, yellow bars; Figure 3C). All *Sfla* Or67b paralogs had strongest responses to ITC compounds (center of the distribution, green bars; Figure 3C). *Sfla* Or67b1 had the narrowest odorant-receptive range, responding to a smaller subset of ITC compounds tested, indicated by the high kurtosis and sharp peak of the tuning curve. Overall, these results demonstrate that *Dmel* Or67b and *Spal* Or67b do not respond to ITCs and have similar odor-response profiles, while each *Sfla* Or67b paralog is differentially responsive to ITCs that are found in diverse mustard species ^49^.

We then tested whether *Sfla* Or67b paralogs differed in their ITC selectivity, possibly allowing flies to differentiate between different mustard plant species. We stimulated OSNs expressing *Sfla* paralogs with serial dilutions of eight selected ITCs that evoked the strongest responses at 1:100 vol/vol (Figure 3B). Because the magnitude of the responses may be reduced when an Or is expressed in at1 OSNs (in comparison with responses when expressed in ab3A OSNs ^63^) we included both non-normalized and normalized median responses for all Or-odorant pairs (Figure 4A-B). In general and as expected, odorant responses increased with increasing odorant concentration. All *Sfla* Or67b paralogs had similar dose-response-curves when stimulated with BITC, while other ITCs elicited Or-specific dose-responses (Figure 4). All three *Sfla* Or67b paralogs are sensitive to ITCs, responding to relatively low odorant concentrations, but have differential ITC selectivity. Lastly, we analyzed the responses of the three *Sfla* Or67b paralogs using principal component analysis (PCA). We found that most ITC responses distributed separately in the odor space when OSNs were tested at 1:100 vol/vol, except for BITC, 4MBITC and 3MPITC, which were separated best at 1:1,000 vol/vol (Figure supplement 4).

**Figure 4.**
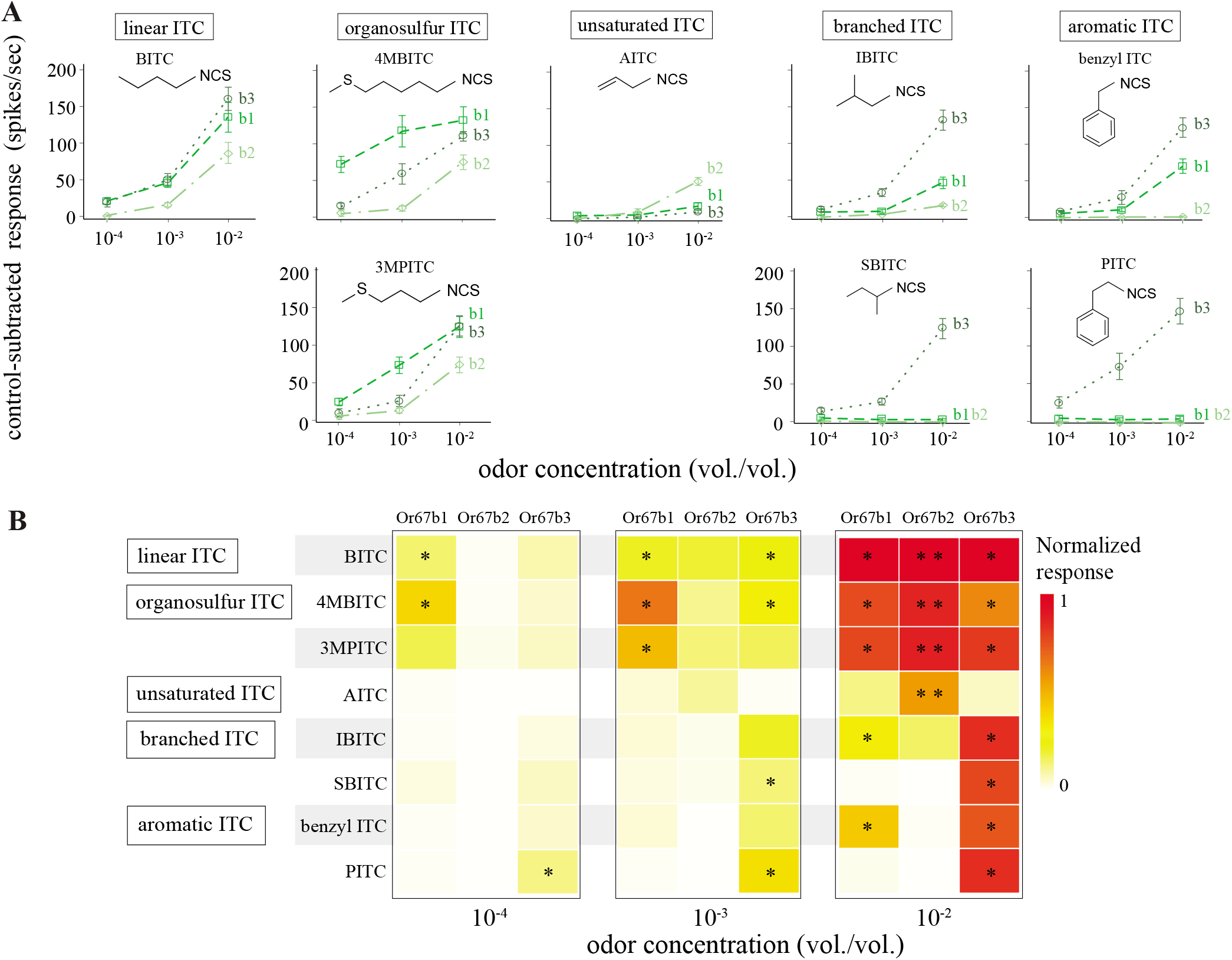
*Sfla* Or67b1-3 have distinct ITC selectivity. **(A)** Dose responses of *Sfla* Or67b1, *Sfla* Or67b2 and *Sfla* Or67b3 (abbreviated as b1, b2 and b3) to stimulation with increasing concentrations (vol/vol) of eight different ITCs (categorized according to molecular structure, top boxes; odorant abbreviations are as in Figure 3). As before, the at1 empty neuron system was only used for expressing *Sfla* Or67b2. Data represent the control-subtracted net number of spikes (average ± SE; n=6-8, obtained from 2-3 animals). **(B)** Heatmap of dose-responses (median, color-coded) from the three *Sfla* paralog normalized (to allow comparisons across paralogs) by each paralog’s median response to 1:100 vol/vol of BITC (the strongest ITC activator across all paralogs). Asterisks indicate significant differences as explained in Materials and Methods (One-sample signed rank tests; * p<0.05; ** p<0.01). The strongest responses were evoked by the highest ITC concentrations, with many compounds evoking responses from all paralogs, particularly in the case of *Sfla* Or67b1 and b3; the number of stimuli that evoked responses decreased with decreasing odorant concentration.

We next tested the extent to which the presence of the ITC functional group (–N=C=S) was necessary for evoking responses from OSNs expressing *Sfla* Or67b paralogs. Therefore, we stimulated OSNs with each of two linkage isomers, BITC (which bears the ITC functional group, –N=C=S) and benzyl thiocyanate BTC (which bears the thiocyanate functional group, -S≡C-N) (Figure 3B, C). Stimulation with BITC produced robust activity in OSNs expressing any of the three paralogs (one-sample signed rank tests, p<0.05), while stimulation with BTC had no effect (p>0.05; Figure 3B, C). This differential activation pattern is therefore likely due to the presence of different functional groups (ITC vs. TC), because these compounds are not only isomers but also have similar volatilities.

### *S. flava* antennal OSNs are responsive to ITCs

*Sfla* Or67b paralogs responded sensitively to diverse ITCs when expressed in the heterologous empty neuron system (Figure 4). We next investigated whether ITC-sensitive Ors are indeed present in the antenna of *S. flava* by recording olfactory responses of *S. flava* antennal basiconic-like (n=36 OSNs) and trichoid-like sensilla (n=36 OSNs: Figure supplement 5) to stimulation with BITC 1:1,000 vol/vol. We used BITC at this low, naturally-occurring concentration, because it evoked responses from all *S. flava* paralogs (Figure 4B; middle panel). We found that 19% of recorded OSNs in basiconic-like and in trichoid-like sensilla showed medium to strong responses (median: 130 and 166 net spikes/second, range: 74-252 and 65-200 net spikes/second respectively in each sensilla type) to stimulation with BITC (Figure supplement 5A, B). These ITC-sensitive OSNs are located proximally in basiconic-like sensilla, and distally in trichoid-like sensilla (Figure supplement 5C). The fact that *S. flava* has BITC-sensitive OSNs in at least two different morphological types of antennal olfactory sensilla comports with our finding that *Sfla* Or67b2 is functional only when expressed in trichoid sensilla of *D. melanogaster*, while *Sfla* Or67b1 and *Sfla* Or67b3 are functional in basiconic sensilla of *D. melanogaster*.

### *S. flava* is attracted to mustard plant odors and volatile ITCs

Because *S. flava* is a mustard plant specialist, and because we showed that *S. flava* Or67b paralogs – but not those from its generalist relatives – respond selectively to ITCs, we hypothesized that *S. flava* has evolved attraction to these odorants. We addressed this using a dual-choice olfactory assay (based on ref. ^64^; Figure 5A; Figure supplement 6) in which flies are allowed to choose between and odor-laden and an odorless arm of a “y-maze” olfactometer ^37^.

**Figure 5.**
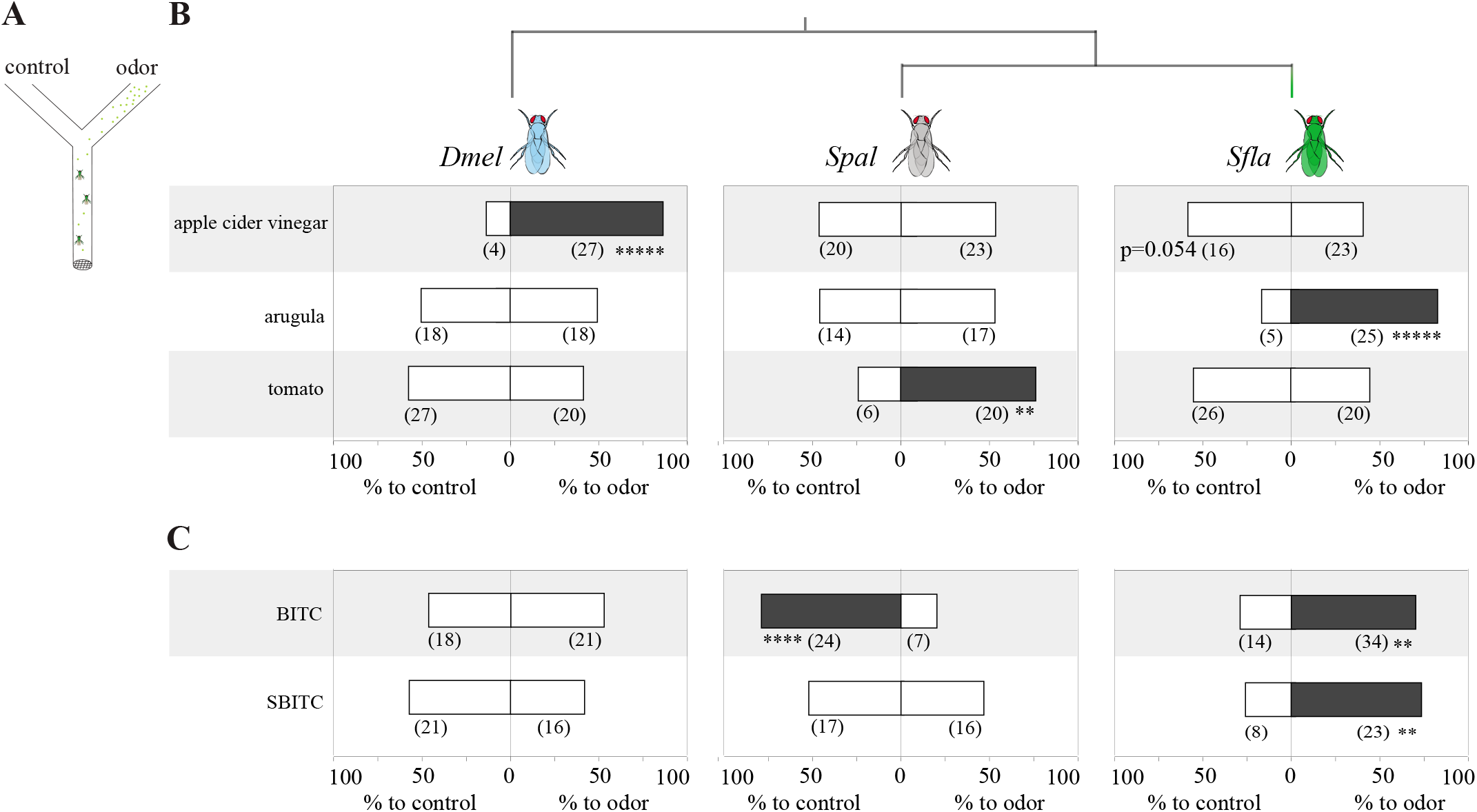
Olfactory behavioral responses of *S. flava* and its microbe-feeding relatives *S. pallida* and *D. melanogaster* to ecologically related odors and ITCs. **(A)** Schematic representation of the dual choice y-maze used to test the olfactory responses of flies (see details in Figure supplement 6, and materials and methods). One arm of the maze offered constant odor/odorant airflow (apple cider vinegar, arugula, tomato or single ITC compounds at 1:100 vol/vol), while the control arm offered a constant odorless airflow (controls: water for odors, and mineral oil for single odorants). In each test a group of non-starved flies (n=3-4) was released at the base of the maze and allowed to choose between the two arms of the maze. Each test (maximum duration=5 min) ended when the first insect (out of all released) made a choice. **(B)** Olfactory behavioral responses of *D. melanogaster*, *S. pallida* and *S. flava* to apple cider vinegar odors and VOCs from leaves of arugula and tomato plants. Data represent the percentage of tests in which animals choose the odorous or odorless (*i.e.* control: water in the case of apple cider vinegar, a piece of tissue paper in the case of plant leaves) arms of the maze; numbers between parentheses indicate the number of tests with choices for one or the other arm. For each fly species and odor/odorant compound, data was analyzed using two-tailed Binomial tests (** p<0.01, **** p<0.001, ***** p<0.0001). *S. flava* was attracted to mustard (arugula) VOCs but not to non-mustard (tomato) VOCs; flies tended to avoid apple cider vinegar odors although differences were not significant (p=0.054); *D. melanogaster* was strongly attracted to apple cider vinegar but not to arugula or tomato leaf VOCs; *S. pallida* was only attracted to tomato leaf VOCs. **(C)** Olfactory behavioral responses of flies from the three species to single ITC compounds (1:100 vol/vol in mineral oil loaded in filter paper); the control arm of the olfactometer offered mineral oil loaded in filter paper. Assays were conducted as described in **B**. *S. flava* was strongly attracted to both ITC compounds tested, *S. pallida* was strongly repelled by BITC, and *D. melanogaster* was indifferent to either ITC.

We first tested the extent to which flies are differentially attracted to mustard and non-mustard leaf VOCs under our experimental conditions. *S. flava* is attracted to arugula and *A. thaliana* leaf VOCs (two-tailed Binomial tests, in all cases p<0.05), while the closely related species *S. pallida* was not (p>0.05; Figure 5 and Figure supplement 7). Leaf VOCs from a non-mustard plant (*e.g.* tomato) did not attract or repel *S. flava* (p>0.05), but attracted *S. pallida* (p<0.01; Figure 5), which comports with the fact that *S. pallida* can be reared in medium containing decaying tomato in the laboratory. *S. flava* tested with leaf VOCs from *A. thaliana* GKO and *PAD3* mutants (which differ in their ability to produce ITCs^50, 52–53^) similarly preferred leaf VOCs from either of these genotypes over clean air (Figure supplement 7). This is not especially surprising because GKO plants still release many VOCs ^50–53^.

As expected, *D. melanogaster* was strongly attracted to apple cider vinegar odors (p<0.05; Figure 5) in agreement with previous studies ^47^. Both *S. flava* and *S. pallida*, in contrast, distributed at random between the apple cider vinegar odor-laden and the odorless arm of the maze (Figure 5). *S. flava* is specifically attracted to mustard plant odors, but not to plant odors in general (represented by those from tomato leaves), and was indifferent to odor sources that attract *D. melanogaster*, characterized by acetic acid and ester, carbonyl, and hydroxyl-containing compounds ^65–66^.

We next investigated whether ITCs alone can mediate olfactory attraction in *S. flava*. We chose BITC and SBITC because these compounds evoked distinct odor responses from *Sfla* Or67b paralogs (Figure 4; BITC strongly activates all *S. flava* paralogs, while SBITC activates only *Sfla* Or67b3). *S. flava* was attracted to BITC and SBITC (p<0.05 in both cases; Figure 5B). Interestingly, *S. pallida* strongly avoided BITC (p<0.005; Figure 5), which must occur *via* an Or67b-independent olfactory pathway because *Spal* Or67b does not respond to any of the ITCs tested, even at high concentrations (Figure 3). Thus, single ITCs compounds, which evoke strong responses from *Sfla* Or67b paralogs and *S. flava* antennal OSNs (Figures 3-4; Figure supplement 5), mediate olfactory orientation in *S. flava* but do not attract (and can even repulse) its microbe-feeding relatives.

### Expression of *Sfla* Or67b in the homologous olfactory circuit of *D. melanogaster* confers behavioral responses to ITCs

Because *Sfla* Or67b paralogs selectively respond to ITCs, we tested if the ectopic expression of these receptors in the homologous olfactory circuit of the microbe-feeder species is sufficient to mediate attraction to these compounds. We first used the Or22a olfactory circuit because it is a circuit known to mediate olfactory attraction in diverse drosophilids ^32–33, 67^. We focused on *Sfla* Or67b3, as this Or has a broader sensitivity to ITC compounds than *Sfla* Or67b1 or *Sfla* Or67b2 (Figures 3 and 4). As before, we used a dual-choice olfactometer and tested flies in which the expression of *Sfla* Or67b3 or *Dmel* Or67b is under the control of GAL4 in the ab3A neuron, as well as the three parental control lines. We tested flies with BITC 1:1,000 vol/vol to avoid compensatory/un-specific responses caused by the lack of Or22a. *D. melanogaster* flies expressing *Sfla* Or67b3, but not *Dmel* Or67b in ab3A OSNs, preferred the BITC-bearing arm of the maze (p<0.05, Figure 6A). Having found that ectopic expression of an Or confers behavioral responses in this set-up, we next expressed *Sfla* Or67b3 in the Or67b-expressing homologous olfactory circuit of *D. melanogaster* (in this experiment flies did not lack expression of the endogenous Or67b). Flies expressing this Or, but not those expressing an additional copy of *Dmel* Or67b (a control), were attracted to BITC (p<0.05; Figure 6B). Ectopic expression of the ITC-responsive *Sfla* Or67b3 in the homologous Or67b circuit of the distantly related (*ca.* 70 million years divergence time) microbe-feeding *D. melanogaster* can confer olfactory responses to ITCs.

**Figure 6.**
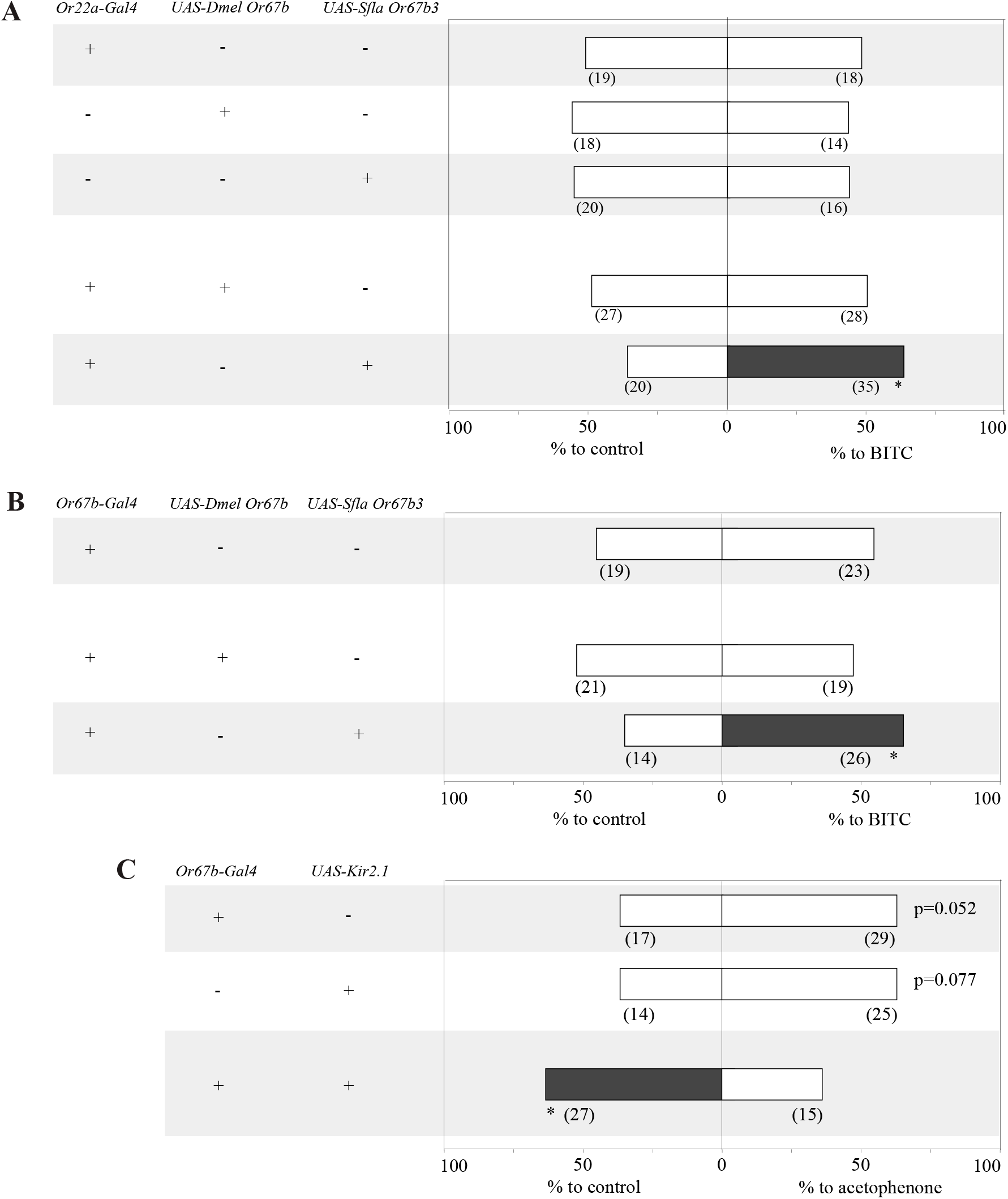
Ectopic expression of *Sfla* Or67b3 in Or22a OSNs or Or67b OSNs conferred behavioral responses to BITC in *D. melanogaster.* **(A)** Behavioral responses of *D. melanogaster* flies expressing *Dmel* Or67b or *Sfla* Or67b3 in Or22a OSNs lacking its cognate olfactory receptor in dual-choice assays (BITC 1:1,000 vol/vol vs. mineral oil control). Experiments were conducted and analyzed as explained in the caption to Figure 5. The two parental control flies (first two groups) and flies expressing *Dmel* Or67b, as expected, were not attracted neither repelled by BITC (Binomial tests, p>0.05 in all cases). In contrast, flies expressing *Sfla* Or67b3 were attracted to the odorant (* p<0.05). These results show that ITCs can evoke olfactory behavioral responses when *Sfla* Or67b3 is expressed in an olfactory circuit that governs attraction. **(B)** Same as **A**, but flies expressed *Dmel* Or67b or *Sfla* Or67b3 in Or67b OSNs (note that flies have the endogenous Or67b expressed in OSNs, in addition to the transgene). As in A, only flies carrying the *S. flava* transgene were attracted to BITC (* p<0.05). **(C)** Behavioral responses of *D. melanogaster* flies expressing a silencer of synaptic activity (Kir2.1) in Or67b OSNs, along with the responses of the two parental control lines (transgenes indicated to the left). One arm of the maze offered acetophenone (1:50,000 vol/vol), a strong *Dmel* Or67b activator ^55^ (Figure 3), while the other arm had mineral oil. Experiments were conducted as explained in the caption to Figure 5, but flies were starved 24 hours previous to testing. As observed for wild-type flies (see Figure supplement 8 and ref. ^68^), genetic control flies showed a trend (0.05<p<0.08) for attraction towards low concentrations of acetophenone, while flies with Or67b OSNs silenced lost attraction and were instead repelled by the odorant (as wild-type flies tested with higher concentrations of acetophenone; see Figure supplement 8 and ref. ^68^). All these findings along with previous reports ^44^ suggest that the ancestral Or67b circuit mediates olfactory attraction.

These results provide an experimental test of the extent to which the Or67b olfactory circuit mediates olfactory attraction in *D. melanogaster*, as has been found for *D. mojavensis* ^44^. To address this, we took advantage of the high sensitivity of *Dmel* Or67b to acetophenone (Figure 3 and ref. ^55^), which evokes behavioral attraction at low concentrations but repellence at high concentrations ^68^. Our experimental conditions faithfully reproduced these results: *D. melanogaster* (Canton-S) was attracted to low concentrations of acetophenone and repelled by high concentrations (Figure supplement 8). Further, we silenced Or67b OSNs using UAS-Kir2.1, an inwardly-rectifying potassium channel that suppresses neuronal activity ^69^ under the control of Or67b-Gal4. Genetic controls showed a tendency for attraction to low concentrations of acetophenone (63% in both cases; p=0.052 and 0.077; Figure 6C), while flies with Or67b OSNs silenced were repelled (Figure 6C; Binomial test, p<0.05). The *D. melanogaster* Or67b olfactory circuit mediates attraction towards low concentrations of a cognate ligand. Aversion to low concentrations of acetophenone in flies with Or67b OSNs inactivated is likely due to activation of other Ors sensitive to this odorant (*e.g.* Or10a ^55^) which mediate odorant repulsion. Activation of OSNs in the *D. melanogaster* Or67b olfactory circuit is necessary for behavioral orientation towards Or67b cognate ligands (*i.e.* acetophenone), whereas this activation is sufficient to confer orientation towards ligands of ectopically expressed Ors (*e.g.* ITCs). Simple evolutionary changes in the odorant tuning of an Or (from acetophenone to ITCs) can change the identity of the odorants (in this case ITCs) that evoke behavioral responses^37–38^. It remains to be investigated whether putative downstream changes in Or67b circuitry^38^ occurred in *S. flava*.

## DISCUSSION

Most herbivorous insect species specialize on a narrow range of host plant species ^70^ that synthesize similar secondary compounds. While these toxins serve to defend against herbivory, ancestrally aversive molecules can become co-opted as attractants to those herbivores that specialize on a particular toxic host plant lineage. We investigated the genetic and functional mechanisms underlying the evolution of attraction to toxic host-plants using *S. flava* as a model. A candidate Or lineage (*Or67b*) was triplicated in a recent ancestor of *S. flava* and experienced rapid protein evolution, resulting in three divergent paralogs (*Sfla Or67b1-b3*; Figure 1). Each *Sfla* Or67b paralogs specifically responded to stimulation with mustard-plant odors and volatile ITCs in heterologous expression systems (Figures 2 and 3) and to a specific subset of ITCs (Figure 4). In contrast, *S. pallida* and *D. melanogaster* Or67b orthologs did not respond to ITCs but showed strong responses to stimulation with apple cider vinegar and a broad range of aldehydes, alcohols and ketones (Figures 2 and 3), consistent with their microbe-feeding niche ^71^. In agreement with these results, recordings from *S. flava* antennal sensilla revealed OSNs sensitive to ITCs (Figure supplement 4). *S. flava*, but not *S. pallida* or *D. melanogaster*, is attracted to volatile ITC compounds (Figure 5). Ectopic expression of *S. flava* Or67b3 in the *D. melanogaster* homologous olfactory circuit conferred odor-oriented behavioral responses to ITCs (Figure 6). Finally, suppression of activity in Or67b positive OSNs in *D. melanogaster* decreased preference to a *Dmel* Or67b cognate ligand, acetophenone (Figure 6). The ancestral Or67b olfactory circuit therefore likely mediates olfactory attraction. Altogether, these results suggest that gene duplication followed by specialization is a mechanism by which specialist herbivores evolve Ors that may mediate olfactory attraction towards ancestrally aversive chemical compounds.

### Evolutionary path of olfactory receptor specialization

Or67b triplication in *S. flava* raises several molecular evolutionary questions, including how this duplication contributed to the generation of novel gene functions. In this regard, several scenarios have been proposed ^22, 72^: (A) neofunctionalization, when one of the duplicated genes (paralogs) acquires a new function after accumulating *de novo* mutations, while the other copy retains ancestral function; (B) subfunctionalization *sensu stricto*, where mutations accumulate in both copies leading to partitioning of ancestral function; and (C) specialization, when subfunctionalization and neofunctionalization evolve simultaneously, yielding gene copies that are functionally different from one another and the ancestral copy ^73–74^. Each *Sfla* Or67b paralog selectively responds to different subsets of ITCs across a range of odorant concentrations, while the proteins from its close relatives did not respond to ITCs (Figures 3 and 4). Or67b evolved new ligand-binding affinities in *S. flava*, indicating a neofunctionalization event. Because each paralog responds to different subsets of ITCs, this shift likely evolved before the duplications of Or67b (Figure 7).

**Figure 7.**
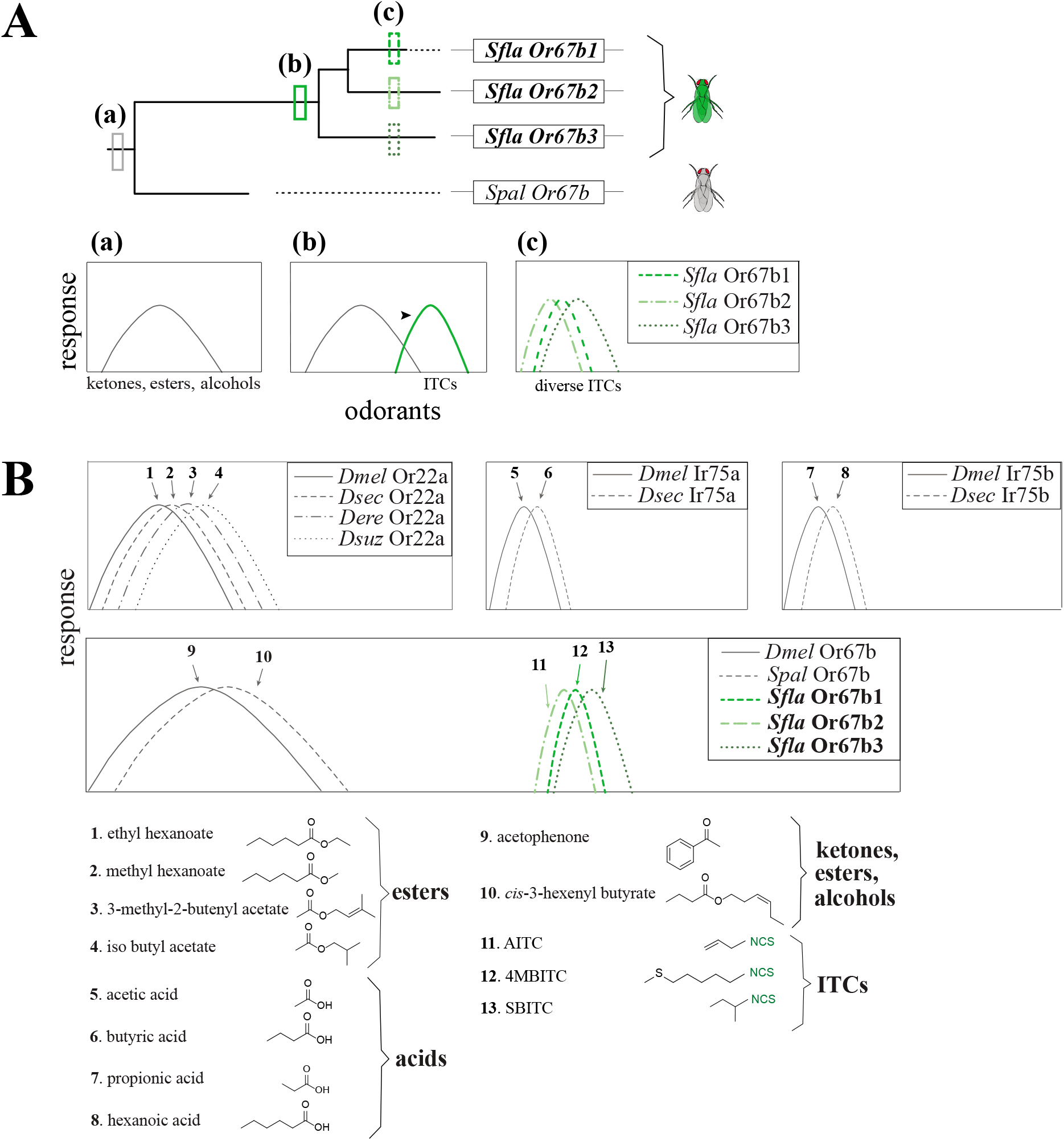
A model for the evolution of *Or67b* and comparison with the known evolution of other Or orthologs in drosophilid flies. **(A)** Model for the evolution of *Scaptomyza Or67b*. The evolution of this Or begins with a shift in the ligand specificity of an ancestral Or67b (a) tuned to *Dmel* activators, GLVs, and ITCs (neofunctionalization, b). Subsequent gene triplication of *Sfla Or67b* gave rise to two additional paralogous *Or67b* genes (*Sfla Or67b1, Sfla Or67b2,* and *Sfla Or67b3*; c), each of them having different but overlapping ITC odorant-receptive ranges (Figures 3 and 4). **(B)** Evolution of drosophilid orthologs with known ligand specificities (Or22a, Ir75a, Ir75b, top; and Or67b, bottom). The Or22a orthologs from *D. melanogaster* (*Dmel* Or22a), *D. sechellia* (*Dsec* Or22a), *D. erecta* (*Der* Or22a), and *D. suzukii* (*Dsuz* Or22a) are all strongly activated by species-specific host-derived esters (compounds 1-4; top left; ^33, 67, 105–106^). The Ir75a orthologs from *D. melanogaster* (*Dmel* Ir75a), and *D. sechellia* (*Dsec* Ir75a) are strongly activated respectively by the acid compounds 5 and 6 ^75^. Similarly, the Ir75b orthologs from *D. melanogaster* (*Dmel* Ir75b), and *D. sechellia* (*Dsec* Ir75b) are respectively activated by the acids 7 and 8 (top right; ^107^). *Dmel* Or67b and *Spal* Or67b are strongly activated respectively by acetophenone and *cis*-3-hexenyl butyrate (compounds 9 and 10), while *Sfla* Or67b paralogs are activated by ITCs only (bottom, paralog-specific activation by compounds 11-13). Note that Or22a, Ir75a and Ir75b orthologs are all divergent but activated by ligands belonging to a single chemical class (whether esters or acids). On the other hand, the ligands of orthologs Or67b from *Dmel* and *Spal* are responsive to a variety of chemical classes which include alcohols, aldehydes and ketones, whereas *Sfla* Or67b orthologs are responsive to ITCs, an entirely different compound chemical class.

### An olfactory receptor sensitive to a new niche-specific chemical class of odorant compounds

Across Drosophilidae, orthologous chemoreceptors respond in a species-specific manner to ecologically relevant ligands. Or22a is a good example of a hot-spot for sensory evolution in *Drosophila* because shifts in ligand binding sensitivity led to different odor-preference behaviors ^37, 67^. Or22a and other highly variable receptors such as Ir75a/b, typically evolve new specificities towards different odorants within one chemical class ^75^ (Figure 7). However, this is not the case for Or67b, which responds to alcohols, aldehydes and ketones in non-herbivorous *D. melanogaster* and *S. pallida*, but in *S. flava*, each paralog is highly sensitive to volatile ITCs, an entirely different chemical class (Figures 3 and 4). The presence of the ITC functional group (–N=C=S) is key to activating these olfactory receptors, highlighting a functional shift in ligand binding specificity (Figure 3B, C). The striking difference in odor selectivity among Or67b orthologs may result from the evolution of herbivory in *Scaptomyza*.

### An olfactory receptor likely mediating attraction to mustard host plants

Through our use of odors produced by crushed leaves of mutant *A. thaliana,* we found that *S. flava* has at least two different olfactory pathways for mustard-plant host attraction: ITC-dependent and ITC-independent (Figure 5 and Figure supplement 7). This is consistent with the fact that plant VOCs are complex bouquets of diverse, lineage-specific molecules that are variably capable of releasing attraction behaviors in specialist herbivores ^76^. Our results do not preclude that other chemoreceptors and OSNs also contribute to mediate olfactory attraction to mustard plant volatiles and ITC compounds in *S. flava*. The necessity of Or67bs for orientation towards ITCs could be probed by generating *S. flava* Or67b loss of function mutants and testing whether flies can still orient towards these odorants. Because *S. flava* females lay eggs in the leaf mesophyll, generation of *Or67b* mutants using CRISPR-Cas9 was not feasible. New technologies such as ReMOT Control ^77^ should enable such experiments in the future. It also remains to be explored if evolution sculpted neural wiring ^37–38^ to modulate responses to ITC compounds.

ITC taste detection in vertebrates and insects leads to aversive behaviors mediated by the contact chemoreceptor TrpA1 ^2, 78–79^. However, volatile ITCs are widely used to trap pests of Brassicaceae ^9^. In agreement with this, the antennae of some of these insects, including *S. flava* ^21^ (Figure supplement 4), respond to volatile ITCs ^10, 80–81^. Similarly, the mustard specialist Diamondback moth *P. xylostella* uses two endogenous Ors selective to ITCs ^10^. These Ors are selective to three ITC compounds, including two (3MPITC and PITC), which strongly activate *S. flava* Or67b paralogs, suggesting that these ITC compounds ^10^ are important for host-orientation in mustard specialists. Importantly, our study further advanced our understanding about the mechanisms underlying evolution of ITC-detecting Ors by comparing Or evolution, function and behavior in herbivorous and non-herbivorous *Scaptomyza* species. The relevance of these gene duplication events is also consistent with the finding that *Or67b* is duplicated only in Brassicaceae-specialist *Scaptomyza* spp.^82^ and not in other herbivorous *Scaptomyza* species specialized on non-Brassicaceae hosts (*i.e. S. graminum*; Figure 1C). Thus, gene duplication and subsequent sequence evolution has played an important role in co-evolution (*sensu lato*) between Brassicaceae plants and diverse herbivorous insects that use them as hosts. More generally, chemosensory specialization in herbivorous insects can result from relatively simple genetic modifications in the peripheral nervous system that change olfactory receptor tuning, contributing to major niche shifts.

## MATERIALS AND METHODS

### Molecular phylogeny of drosophilid olfactory Olfactory Receptors (Or)

Translations of *Ors* from *D. grimshawi*, *D. mojavensis, D. virilis* and *D. melanogaster* (builds dgri r1.3, dmoj r1.3, dvir r1.07 and dmel r6.28, respectively) were downloaded from Flybase (www.flybase.org; ^83^). *S. flava Or* sequences were previously published ^21^. A total of 309 sequences were aligned in MAFFT v7.017 with the E-INS-I algorithm and then manually adjusted ^84^. Models were fitted to the alignment using IQ-Tree and tested using the cAIC criterion ^85^. A maximum likelihood (ML) phylogeny was generated using the Or protein alignment (in RAxML v8.2.10) with the CAT model of rate heterogeneity with seven distinct categories, the JTT substitution matrix, empirical amino acid frequencies, and 1,000 rapid bootstraps ^86^. Orco sequences were designated as the outgroup as is standard practice in evolutionary analyses of arthropod Ors.

### Time calibrated molecular phylogeny of *Scaptomyza* and *Drosophila* spp

A time-calibrated species tree was inferred using the loci: *16S*, *COI*, *COII*, *ND2*, *28S*, *Cad-r*, *Gpdh1*, *Gstd1*, *Marf*, *l(2)tid* (also known as *Alg3* or *Nltdi*) and *AdhR* from 13 spp of *Drosophila* and four *Scaptomyza* spp. *Scaptomyza* sequences were accessed from the genomes in refs. ^13, 87^ using tblastn searches for protein coding genes and blastn searches for the ribosomal RNA genes. The *Adh Related* gene appears to be deleted in *S. flava* and was coded as missing data. Two uniform priors on the age of *Scaptomyza* and the age of the combined virilis-repleta radiation, Hawaiian *Drosophila* and *Scaptomyza* clade were set as in ref. ^88^. Protein coding genes were portioned into first and second combined and third position partitions, except *COI and Gpdh1*, which were divided into first, second and third partitions. The partitioning scheme was chosen based on Partition Finder 2.1.1 ^89^ and nucleotide substitution models chosen with IQ-Tree ^85^. Model parameters, posterior probabilities, accession numbers and genome coordinates can be found in table 1. Five independent runs of BEAST v2.6.2 ^90^ each with 25 million generations were run logging after every 2500^th^ generation to infer the chronogram with 10% burn-in. Phylogenies were also inferred in RAxML v8.2.12 ^86^ with the GTR+GAMMA+I model and 1000 rapid bootstraps and with parsimony in PAUP* 4.0a ^91^ with TBR branch swapping and 1000 bootstraps. Phylogeny parameters, sequence accession numbers and likelihood scores available in supplementary dataset 1.

### Molecular phylogeny of drosophilid *Or67b* genes

*Or67b* coding sequences (CDS) from *D. grimshawi*, *D. mojavensis, D. virilis D. sechellia, D. simulans*, *D. erecta, D. yakuba, D. pseudoobscura, D. persimilis, D. ananassae* and *D. melanogaster* (builds dgri r1.3, dmoj r1.3, dvir r1.07, dsec r1.3, dsim r1.4, dere r1.3, dyak r1.3, dpse r3.2, dper r1.3, dana r1.3 and dmel r6.28, respectively) were downloaded from Flybase (www.flybase.org; ^83^) and *D. hydei* from Genbank (accession number XM_023314350.2). The *S. pallida* DNA sequence was obtained through PCR and Sanger sequencing as described below. *S. flava* DNA sequences were previously published ^21^. Two more *Scaptomyza* sequences were obtained from refs. ^13, 87^ including the non-leaf-mining species *S. hsui* (subgenus Hemiscaptomyza and a microbe-feeder) and *S. graminum* (subgenus *Scaptomyza* and a leaf-miner on Caryophyllaceae). DNA sequences were aligned, partitioned by codon position, models fitted to all three partitions and chosen according to AICc (GTR+I+G) in IQ-Tree ^85^. Trees were inferred using RAxML (v8.2.10) with the GTRGAMMA+I model and 1000 rapid bootstraps, and MrBayes (v3.2.6) setting Nst to 6, nucmodel to 4by4, rates to Invgamma, number of generations to 125,000, burnin equal to 20% of generations, heating to 0.2, number of chains to 4, runs to 2 and priors set to default setting ^92^. An additional parsimony analysis was performed in Paup 4.0 ^91^ with TBR branchswapping and 1000 bootstraps. Model parameters, accession numbers and likelihood scores available in Supplementary file 1.

### Analysis of molecular evolutionary rates

CDS of homologs of every *Or* gene in *S. flava* found in the twelve *Drosophila* genome builds were aligned to *S. flava Or* CDS. Homology was assessed according to inclusion in well-supported clades in the *Or* translation phylogeny from the twelve species; *S. flava* sequences were previously published ^21^. Sequences were aligned in MAFFT (v7.017) ^84^ and adjusted manually to preserve codon alignments. *Or98a*-like genes found in subgenus *Drosophila* species were split into three separate clades, as were a group of *Or83c* paralogs not found in *D. melanogaster*, and a group of *Or85a-*like genes. All examined sequences of *Or46a* contain two alternatively spliced exons, so this gene was analyzed with all gene exon sequences in a single alignment. *Or69a*, however, contains alternatively spliced exons only in species within the subgenus *Sophophora*, and therefore these alternative splice forms were analyzed as separate taxa. Phylogenies were generated for every alignment using PhyML ^93^ with the GTR+G substitution models. If >70% bootstrap support was found for a topology contrary to the known species topology, or if the *Or* homology group contained duplicates, these trees were used in PAML analyses instead of the species tree.

Our next goal was to identify *Or* genes experiencing rapid rates of protein evolution. Branch models of sequence evolution were fit using PAML 4.9h ^41^. A foreground/background branch model was fit for every *S. flava* tip branch and every ancestral branch in a *Scaptomyza*-specific *Or* gene duplication clade, and compared in a likelihood ratio test to a null model with one d*_N_*/d*_S_* rate (ratio of non-synonymous to synonymous substitution rate) for every unique phylogeny (75 tests in total). After focusing on *Or67b*, patterns of molecular evolution among the drosophilid *Or67b* homologs were explored using the expanded *Or67b* CDS phylogeny above. Foreground/background branch models were fit for every branch in the *Or67b* phylogeny and the *S. flava* Or67b paralogs clade with likely ratio tests performed as above (34 tests total, table 2). P-values were adjusted for multiple comparisons using the false-discovery rate (FDR) method ^94^. Branch test results and model parameters in Supplementary file 2.

### Synteny analysis between S. flava, S. pallida and D. melanogaster Or67b scaffolds

Five genes up and downstream of each *Or67b* ortholog in *S. flava* were extracted using annotations from a current GenBank assembly (GenBank Assembly ID GCA_003952975.1), shown in Supplementary file 2. These genes are respectively known as pGOIs (genes proximal to the gene of interest, or GOI). To identify pGOIs, we used tBLAST to the *D. grimshawi* genome, the species closest to *S. flava* with a published annotated genome (GenBank Assembly ID GCA_000005155.1). By identifying the pGOI scaffolds, we determined that there was only one copy of *Or67b* in *D. grimshawi*, which is syntenic with the *S. flava Or67b2* ortholog. We determined that this copy is also syntenic with the single *Or67b* copy of *S. pallida* (Figure supplement 2) and *D. melanogaster*.

### Fly husbandry and lines

*D. melanogaster* (wild-type Canton-S and transgenic lines) were reared using standard cornmeal media, yeast, and agar medium prepared by UC-Berkeley core facilities. Isofemale lines of *S. pallida* (collected in Berkeley, California, US) were maintained on Nutri-Fly medium (Genesee Scientific). *S. flava* (collected in New Hampshire, US) were maintained on fresh *A. thaliana* plants and 10% honey water solution. All flies were cultured at 23°C and 60% relative humidity under a 12-h light/12-h dark cycle. *S. flava* and *S. pallida* were ca. 7-10 days old at the time of experiments; *D. melanogaster* (wild-type or transgenic) were ca. 3-10 days old at the time of the experiments.

Flies used for heterologous expression of Ors were of the following genotypes: for flies with ab3a “empty neuron” system, *Or22ab^Gal4::3xP3-DsRed^* was used. This “M2-MD” line was generated by a CRISPR-Cas9 mediated deletion of *Or22a/b* and a knock-in of *Gal4* and *DsRed* ^95^ by homology directed repair (HDR); in these flies *Gal4* is not functional. Therefore, we used *Or22ab^Gal4::3xP3-DsRed^; Or22a-Gal4/UAS-Or67b* flies were used for experiments. For flies with the at1 “empty neuron” system, *Or67d^Gal4^* line ^28^ was used. The functional absence of *Or22a* and *Or22b* genes in M2-MD flies, or functional absence of *Or67d* in the *Or67d*^GAL4^ line, were respectively confirmed by electrophysiological analysis on ab3A neurons or at1 neurons (Figure supplement 3). The M2-MD line was used to generate flies expressing *Dmel Or67b*, *Spal Or67b*, *Sfla Or67b1* and *Sfla Or67b3* under the control of *Gal4* in the ab3A “empty neuron” ^46^. Similarly, the *Or67d*^GAL4^ line was used to generate flies expressing *Sfla Or67b2* under the control of Gal4 in the at1 “empty neuron” ^28^. The progeny of those crosses was then used for single sensillum recordings and in some cases for behavioral assays. The *UAS-SflaOr67b1*, *UAS-Sfla Or67b2*, *UAS-Sfla Or67b3*, and *UAS-Spal Or67b* strains were generated during this study. The *Or67b-Gal4* fly line (BDSC# 9996) was crossed with *UAS-Or67b* lines or *UAS-Kir2.1* ^69^ for behavioral assays.

### *Scaptomyza Or67b* gene cloning, UAS line generation, and verification of *S. flava Or67b* transcription

The *UAS-Or67b* transgene lines were constructed as follows: RNA was extracted from 20–25 days post-emergence adults of both sexes from laboratory-reared *S. pallida* (collected from the White Mountains, New Mexico, USA) and *S. flava* (collected from near Portsmouth, New Hampshire, USA). RNA was extracted using Trizol (Thermo-Fisher, Waltham, MA, USA) and precipitated with isopropanol. Extracted RNA was treated with DNaseI, cDNA was generated using qScript cDNA Supermix (Quantabio, Beverly, MA, USA). Absence of genomic DNA in cDNA preparations was verified by attempting to PCR-amplify fragments of the *Marf* gene from reactions lacking reverse transcriptase (Source data). PCR conditions and primers are detailed in ^12^. CDS plus 7–9 bp of untranslated sequence were amplified using High Fidelity Phusion Taq (New England BioLabs, NEB, USA), 3% DMSO vol/vol, and the PCR primers (Supplementary file 4) with the following program: initial denaturing at 98°C during 30 sec; 35 cycles at 98°C during 10 sec, 58°C during 30 sec, 72°C during 45 sec, and extension at 72°C during 7 min. PCR fragments of the expected size were purified using the Qiaquick Gel purification kit protocol (Qiagen). An overhang was added to purified *Or67b* amplicons with Taq polymerase (Fermentas) and cloned using the pGEM-T Easy cloning kit protocol (Promega). Plasmids were extracted and purified using the GenElute plasmid miniprep kit (Sigma-Aldrich, St. Louis, MO, USA). EcoRI and KpnI cut sites were introduced using restriction enzyme cut-site primers (Supplementary file 3) with 10 ng/µL diluted plasmids (as template) with 3% DMSO vol/vol and the following program: initial denaturing at 98°C during 30 sec; 35 repetitions of 98°C during 10 sec, 55°C during 50 sec; 72°C during 45 sec; and final extension at 72°C during 7 min. The pUAST attB plasmid ^96^ and the four *S. pallida* and *S. flava Or67b* PCR amplicons with RE flanking sites were individually double-digested with KpnI and EcoRI high-fidelity enzyme in cut smart buffer for 3 hours, according to the manufacturer’s protocol (NEB). Cut fragments were gel-purified using the Qiaquick Gel Cleanup Kit (Qiagen) and ligated in a 1:3 vector:insert molar ratio using T4 ligase (Promega). Ligations were transformed into JM109 cells. Some cells were preserved as glycerol stocks and a portion were used for injection into the y^1^ w^67c^^23^; P{CaryP}attP2 *D. melanogaster* line for generating *Sfla Or67b1, Sfla Or67b2, Sfla Or67b3, Spal Or67b* or into the y^1^ w^67c23^; P{CaryP}attP40 for *Sfla Or67b2* (BestGene Inc., Houston, Texas, USA). Transformants were selected from individually injected flies with compound eye color rescue phenotypes.

### Single sensillum recordings (SSR)

All *Or67b* constructs were expressed in the ab3A empty neuron system, except *Sfla Or67b2*, which was expressed in at1 empty neuron system because this Or exhibited spontaneous activity only in at1 trichoid OSNs in transgenic *D. melanogaster* flies (Figure supplement 3).

Fed, adult female flies were prepared for SSR as previously described ^45^. We identified the antennal sensilla housing the OSN(s) of interest using an Olympus BX51WI upright microscope with 10x and 50x objectives (Olympus, UPlanFL N 10x, UplanFL N 50x). We recorded the responses of 6-10 sensilla obtained from 2-4 individuals for each experiment/odorant, the standard in this type of experiment (*e.g.* ref. ^97^). Extracellular activity was recorded by inserting a tungsten electrode into the base of either ab3 or at1 sensilla. Signals were amplified 100x (A-M systems, Differential AC Amplifier model 1700), digitized using a 16-bit analog-digital converter, filtered (low cut-off: 300 Hz, high cut off: 500 Hz), and analyzed off-line using WinEDR (v3.9.1; University of Strathclyde, Glasgow). A tube delivering a constant flow of charcoal-filtered air (16 ml/min, using a flowmeter; Gilmont instruments, USA) was placed near the fly’s head, and the tip of the stimulation pipette (50 ml) was inserted into the constant air stream. The stimulation pipette contained a piece of filter paper loaded with 20 µl of odorant solution or the solvent control. One second pulse of clean air was delivered to the stimulus pipette using a membrane pump operated by a Stimulus Controller CS 55 (Syntech, Germany). Ab3 sensilla were identified by using three standard diagnostic odorants ^98^ (all odorants were obtained from Sigma-Aldrich, US, purity > 95%): ethyl hexanoate (CAS # 123-66-0), ethyl acetate (CAS # 141-78-6) and 2-heptanone (CAS # 110-43-0) (Figure supplement 3). We were able to distinguish at1 sensilla from at2 and at3 because the former houses a OSN, whereas at2, at4 and at3 respectively house two, three and three OSNs ^98^.

The following odor sources (purchased from Berkeley Bowl in Berkeley, California, USA, unless otherwise mentioned; 20 µl of material were loaded on filter paper unless noted) were used: apple cider vinegar (40 µl, O Organics, USA), grated roots of the four Brassicaceae species; *W. japonica* (wasabi), *A. rusticana* (horseradish), *B. rapa* (turnip), *R. sativus* (daikon), and the Amaranthaceae species *B. vulgaris* (beet). Approximately 10 g of roots were grated immediately before experiments and used right away to prevent compound degradation. Brassicaceae species *E. vesicaria* (arugula), *A. thaliana*, and a Solanaceae species, *S. lycopersicum* (tomato), were grown from seeds at 23°C and 60% relative humidity under a 12-hour light: 12-hour dark cycle, and leaves from 3-8 weeks old plants were used for odor stimulation. The following *A. thaliana* genotypes were used: wild-type (Col-0), glucosinolate knockout (GKO) mutant in *myb28 myb29 cyp79b2 cyp79b3*, which has no detectable aliphatic and indolic glucosinolates nor camalexin ^50, 52–53^, and camalexin-deficient *phytoalexin deficient 3* (*PAD3*) mutants that have wild-type levels of aliphatic and indolic glucosinolates (but no camalexin). Therefore, *PAD3* plants and are more appropriate controls for comparisons with GKO plants than Col-0 plants ^54^. Three-four leaves were excised from plants and homogenized with a grater immediately before tests; the homogenate was replaced every 30 min since OSN odor responses were stable at least during this time window. All synthetic odorants were diluted in mineral oil (1:100 vol/vol) unless otherwise noted. The following odorants (all from Sigma-Aldrich, US, purity >95%) were diluted in dimethyl sulfoxide (DMSO): mandelonitrile (CAS # 532-28-5), iodoacetamide (CAS # 144-48-9), N-methylmaleimide (CAS # 930-88-1), N-hydroxysuccinimide (CAS # 6066-82-6), and benzyl thiocyanate (CAS # 3012-37-1). 11-*cis* vaccenyl acetate (CAS # 6186-98-7) and 4MBITC (CAS # 4430-36-8) were diluted in ethanol. All chemicals used in this study are listed in supplementary file 6.

The “net number of spikes” was obtained by counting the number of spikes originating from the OSN of interest during a 1-second window which started 0.2 seconds after the onset of stimulation, and subtracting from this number the background spiking activity (obtained by counting the number of spikes in a 1-second window prior to the onset of the odor stimulation). In all figures (unless otherwise noted) the net response to each odor or odorant stimulation was subtracted from the median net response to the solvent control used (some odorants were dissolved in solvents other than mineral oil, see preceding paragraph).

In SSR experiments it is common to observe unspecific, slight increases/decreases in spiking activity which are not considered biologically meaningful, including small responses to solvent controls. Accordingly, it is not uncommon to find reports, including investigations using the empty neuron system of *D. melanogaster* ^97^, in which net spiking responses smaller than 10-25 spikes/second are not considered true odor-evoked responses (*e.g.* ^99–100^). Therefore, we asked if the net responses to a given combination of Or and odorant compound are statistically significant (p<0.05) using one-sample signed rank tests under the following null and alternative hypotheses:

*Ho:* net number of spikes >-10 and <10

*Ha:* net number of spikes <-10 or >10

Data were also analyzed using Mann-Whitney U tests for comparing two independent groups, Kruskal-Wallis ANOVAs for comparing more than two independent groups (followed by post-hoc tests if significant), and Wilcoxon-matched pairs tests for comparing two paired groups. Although results were considered significant only if p<0.05, we indicated cases in which p-values were slightly larger (0.05<p<0.08). We informed these values because “a result that does not meet the p<0.05 threshold should not be considered meaningless” ^101^ (specially with low sample sizes due to the nature of the experiment, as in this report) “when in fact provides…at least preliminary evidence that requires further attention” ^101^.

Tuning curves and kurtosis values were generated and calculated in Microsoft Excel (2016). Similarly, a matrix of median responses (control-subtracted) was produced and used for PCA in R statistical software. For generation of the heatmap in R (Figure 4B), the median responses of OR-odorant pairs were normalized to the maximum median response (BITC at 1:100 vol/vol) for each Or across all odorants. This normalization served to adjust for the potential intrinsic differences in response magnitude between the ab3A and at1 empty neuron systems ^28^.

### Behavioral tests

The olfactory responses of mated, fed adult female *D. melanogaster* (Canton-S), *S. pallida*, *S. flava,* and transgenic flies were tested using a custom-made dual-choice “Y-shaped” olfactometer ^64^ (Figure supplement 6). Flies were starved 24 hours for experiments shown in Figure 5-6. The “Y piece” of the olfactometer was a propylene connector, and the arms of the “Y” were each connected to a 1-ml syringe containing a piece of filter paper (6 x 50 mm) loaded with the odor or control stimuli. Charcoal-filtered air was delivered to each of the two stimulus syringes using silicon tubing at 250 ml/min; thus, at the base of the maze the air flow was approximately 500 ml/min. Two hours (in the case of *D. melanogaster* and *S. pallida*) or *ca*. 20 hours before tests (in the case of *S. flava*) insects were gently anesthetized under CO2 and placed in groups of thee-four in open-top and mesh-bottom cylindrical release containers (20 mm long x 10 mm diameter) constructed using silicon tubing. The open top of the containers was capped with a piece of cotton soaked in distilled water (in the case of *D. melanogaster* and *S. pallida*) or with a piece of cotton soaked in 10% vol/vol aqueous honey solution (in the case of *S. flava*). Before tests, each release tube was placed on ice for 45-60 seconds to slow down insect activity; the cotton cap was then removed and the open-top of the tube was carefully slid into the open end of the Y maze. Thus, upon being released, insects could walk upwind towards the “decision point” (intersection of the short and long arms of the “Y”) and turn towards either the odor-laden or the odorless arm of the maze. Although three-four insects were released at once (to increase experimental efficacy), only the first choice (and the time of the choice) was recorded; a choice was considered as such only if the insect walked past at least 10 mm into one of the arms, orienting upwind. The test was discarded if two or more insects chose different arms of the maze within a five-second window of each other. Each test lasted a maximum of five minutes, and each group of insects was used only once. Test stimuli were randomly assigned to flies prepared for behavioral tests. As much as possible, insects from the same cohort were tested in the same day with different odors/odorants. In the case of experiments using transgenic fly lines and the progeny of crosses between them, we conducted experiments with the progeny of at least 4-5 independent crosses; control and test flies were tested in parallel as much as possible. Tests with each combination of fly line (or species) and stimulus were conducted in at least five different days with different progeny to compensate for possible day-to-day and cohort variations.

In general, results from an individual test session with a given odor/odorant were discarded if insects did not make a choice in more than 50% of tests (this happened in less than 5-10% of experimental sessions except for *S. pallida*, which often had low activity levels), as it is the standard in this type of experiment. The position of the odor and odorless arms was switched every 1-2 tests to control for positional asymmetries; the mazes and odor sources were changed and replaced for clean/new ones every 4 tests or 10 minutes, whichever occurred first.

The odor/odorant was loaded on a piece of filter paper and inserted into the 1 ml syringe immediately prior to each test; control syringes had a piece of filter paper loaded with the mineral oil solvent (for odorant solutions) or water (in tests apple cider vinegar). Experimental and control filter papers were replaced by fresh ones every 4 tests or about 10-11 minutes, whichever came first. The odorants (20 µl of 1:100 vol/vol mineral oil solution unless noted) used in experiments were BITC (Sigma-Aldrich, CAS # 592-82-5, USA) and SBITC (Sigma-Aldrich, CAS # 15585-98-5, USA), and acetophenone (Sigma-Aldrich, CAS # 98-86-2, USA) in one experiment (Figure 6C). We also used apple cider vinegar (40 µl, O Organics, USA; 40 µl of distilled water was a control stimulus in these tests). For tests of host-orientation, leaves from four to six weeks old *E. sativa*, *A. thaliana,* or *S. lycopersicum* plants, grown in an insect and insecticide/pesticide free chamber or greenhouse, were excised and gently broken just before tests and placed in 5 ml syringes connected to the Y-maze; control syringes had two pieces of wet tissue paper. Plant material and stimulation syringes were replaced by new ones every four tests (*e.g.* after a potential maximum of 20 minutes since excision from the plant, but more often after only 8-10 minutes). In all cases the Y-mazes, tubing and syringes were washed with 70% ethanol and allowed to air-dry before reusing. Experiments were conducted during the 2^nd^-5^th^ hour of the insects’ photophase at 24 °C under white light (Feit electric, 100 Watts; in the case of *S. pallida* and *D. melanogaster*). Green light (Sunlite green, 100 Watts) was used in the case of experiments with *S. flava*, as anecdotal observations suggest that this fly species is more active under this light color, itself a potentially biologically meaningful response. In all cases the total number of tests in which at least one insect chooses one or the other arm of the maze is indicated in the figures for each species/fly line/odor. For behavioral tests, we established *“a priori”* a minimum sample size (n=30); sample sizes for experiments with *S. pallida*, which is unusually inactive, were sometimes lower but averaged n=28 across experimental series (Figure 5).

Because *S. flava* females lay eggs in leaves, generation of mutants using CRISPR-Cas9 was not possible at the time of these experiments. Therefore, we conducted a gain of function tests in which *D. melanogaster* flies expressed *Sfla* Or67b3 in the basiconic ab3A “empty neuron”, or in ab9b OSNs (which express Or67b in *D. melanogaster* ^102^). Controls and experimental flies were tested with butyl-ITC 1:1,000 vol/vol. To investigate the role of the Or67b circuit in mediating olfactory attraction, we expressed UAS-Kir2.1, a genetically encoded inwardly rectifying potassium channel that by keeping cells hyperpolarized prevents neuronal excitation ^69^, under the control of Or67b-Gal4. Experimental and control fly lines were starved during 24 hours (but provided with a wet tissue paper) and tested with acetophenone 1:50,000 vol/vol. To verify that previously reported behavioral responses to acetophenone ^68^ occurred under our experimental conditions, *D. melanogaster* flies (Canton-S) were also tested with various concentrations of this odorant (Figure supplement 8).

For each odor/odorant, species, and fly line, the number of tests in which an insect made a choice for one or the other arm of the Y-maze (with the criteria described above) were tested against a 50% expected random distribution using two-tailed Binomial tests ^103^. Results were considered statistically significant if p<0.05. Data collection for each experimental series ended when significance was achieved, or when n=55, regardless of the outcome. This criterion was adopted because binomial tests require an untenably large sample size to evince small deviations from an expected proportion (*e.g.* 78 tests are required to detect a 10% deviation from the expected 50% random choice at the p<0.05 level). In cases where n<55, we conducted a similar number of tests for control and experimental lines to ensure that results are fairly comparable. Similarly, we also noted behavioral results when p-values were >0.05 but <0.08, because such outcomes indicate that significance (p<0.05) could likely be achieved by increasing sample size ^101^ (although increasing sample size is often difficult for behavioral experiments). For instance, a 10% deviation from the expected 50% distribution and n=55 yields p=0.08, while that same deviation requires n≥74 for achieving p<0.05. Thus, p-values slightly larger than 0.05 bear relevant information because they are indicative of trends that might become significant if sample size were increased. Statistical power was in most significant cases >0.8; exceptions include cases where deviations from the expected 50% distribution were relatively small (*e.g.* a significant 10% deviation requires n=188 to achieve 0.8 statistical power).

### Data Analysis and figure generation

All images and drawings are originals prepared by the authors. Figures were prepared via a combination of WinEDR (v3.9.1), R Studio (v1.2.1335), Microsoft Excel (2016), Adobe Illustrator (2019), ChemDraw (19.0), Python, and Geneious (10.0.9).

## Acknowledgments

We are grateful to Drs. Chauda Sebastian, Dennis Mathew, John Carlson, Barry J. Dickson and Bloomington Drosophila Stock Center (NIH P40OD018537) for sharing *M2-MD*, *UAS-Dmel Or67b,* and *Or67d^Gal4^*, and to Drs. Johannes Bischof and Konrad Basler for donation of the pUASTattB plasmid. C.E.R. thanks Dr. Kristin Scott for support and encouragement. T.M. thanks Dr. Makoto Hiroi for advice on SSR experiments. We thank members of the Whiteman and Scott Laboratories for discussions and comments on the manuscript. This work was supported by the Uehara Memorial Foundation (award number 201931028 to T.M.), the National Institute of General Medical Sciences of the National Institutes of Health (award number R35GM119816 to N.K.W.) and the National Science Foundation (award number IOS 1755188 and DEB 1601355 supporting B.G.-H.).

---

**Figure supplement 1.**
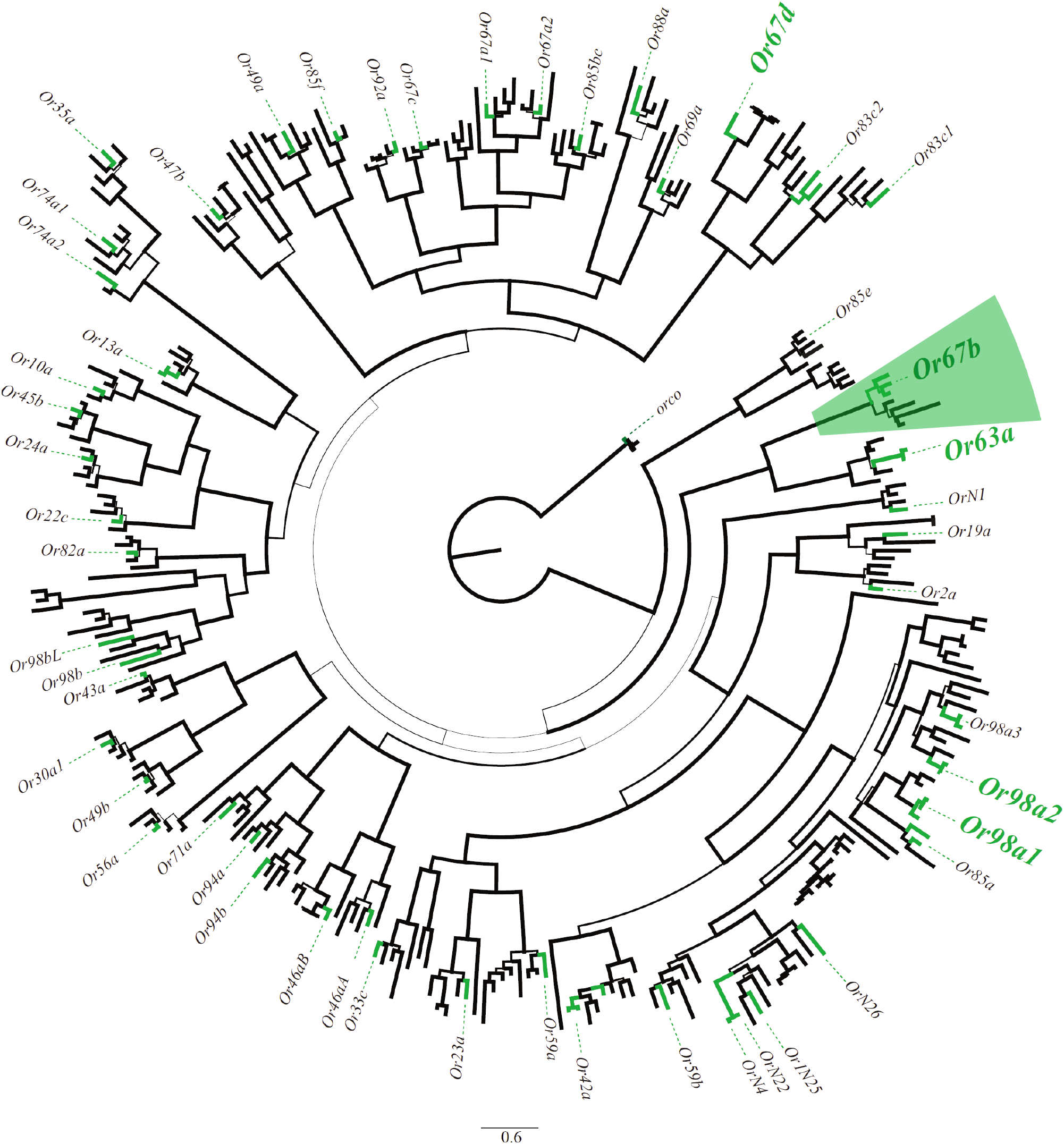
Maximum likelihood (ML) phylogeny of *Ors* in Drosophilidae. ML phylogeny reconstructed from protein translations of the *Ors* found in *S. flava*, *D. melanogaster*, *D. grimshawi*, *D. virilis* and *D. mojavensis* genomes. Line width of branches are proportional to bootstrap support. Green branches indicate *Scaptomyza Ors*. Enlarged gene names in bold include branches with estimated d*_N_*/d*_S_* >1. Scale bar units are substitutions per site.

**Figure supplement 2.**
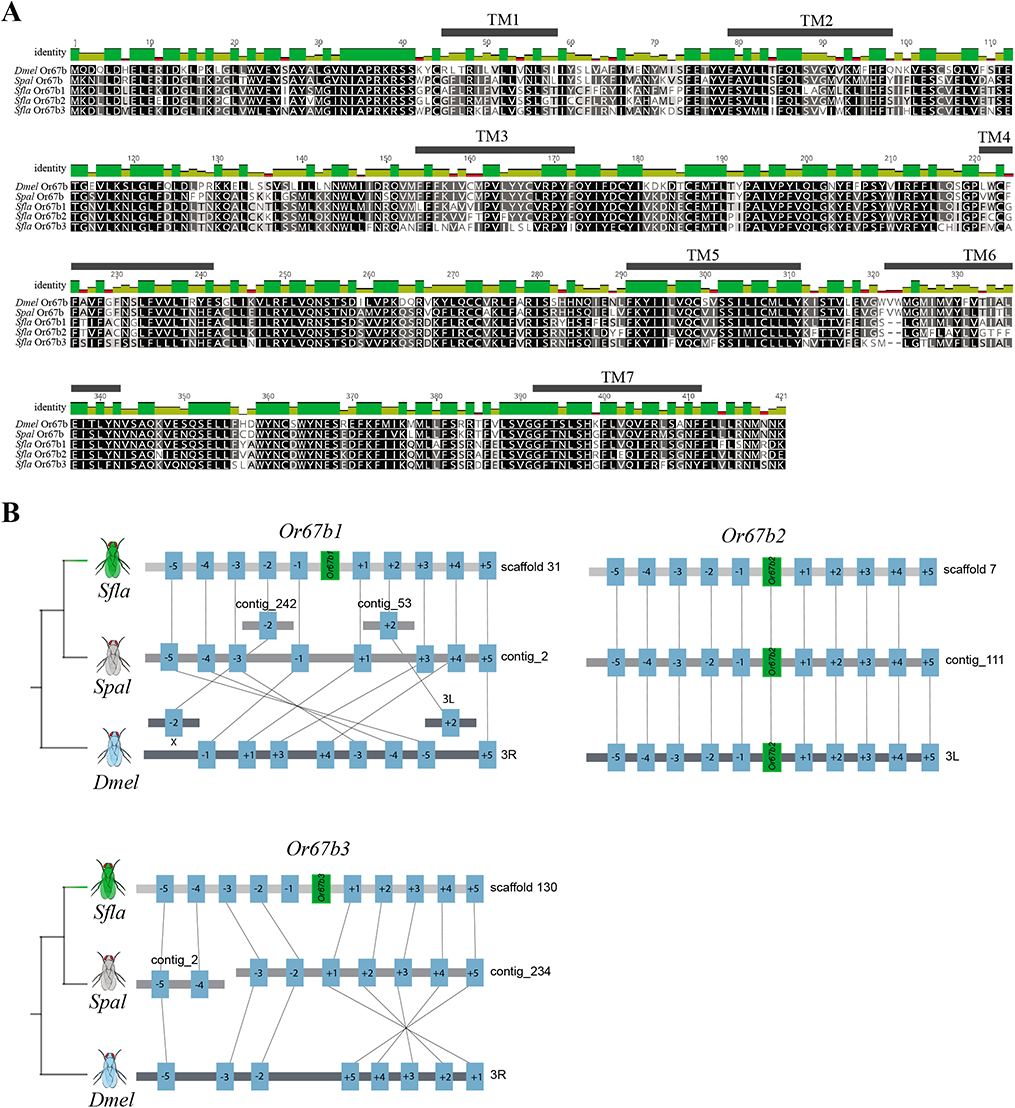
Or67b protein alignment and micro-syntenic patterns of scaffolds from *S. flava, S. pallida*, and *D. melanogaster.* **(A)** Alignments of Or67b proteins. Note that orthologs and paralogs share a large number of amino acids (black squares). Darker colors illustrate higher degrees of sequence similarity, and lighter colors denote residues with high variability in sequence across paralogs and orthologs. Also shown are each of the seven predicted transmembrane domains (TM1-7). **(B)** Micro-syntenic patterns of Or67b scaffolds. Five genes up and downstream of each *S. flava Or67b* ortholog are shown. We determined that there was only one copy of *Or67b* in both *S. pallida* and *D. melanogaster*. Note that in both *S. pallida* and *D. melanogaster*, many of the pGOIs are still present and syntenic with those in *S. flava,* suggesting that the absence of the other *Or67b* orthologs is not a consequence of mis-assembly but a *bona fide* absence.

**Figure supplement 3.**
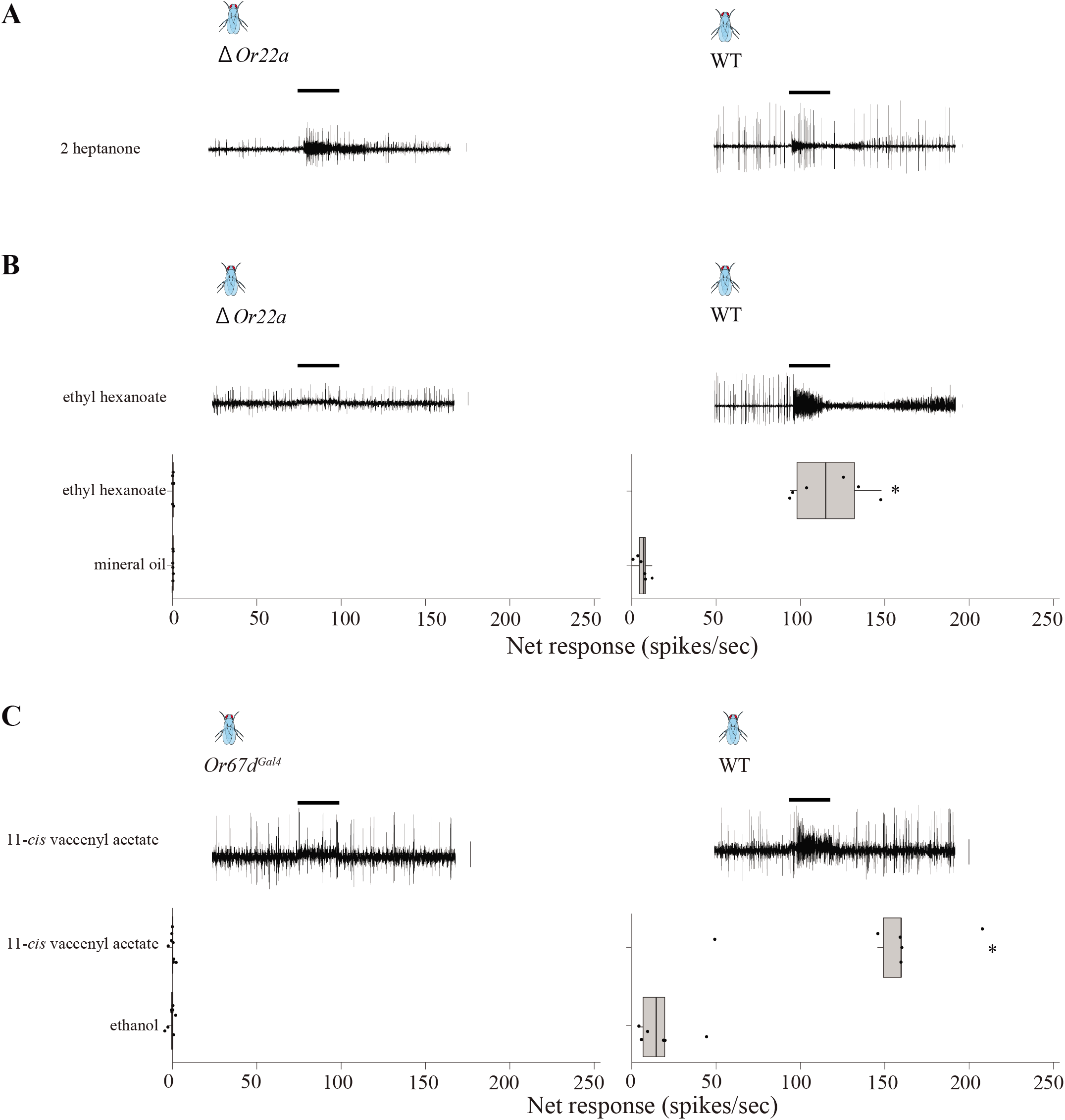
Or22a and Or67b are not expressed in the ab3A and the at1 empty neuron systems. **(A)** Recordings from ab3 sensilla in empty neuron mutants (left) and wild-type flies (right). Representative electrophysiological traces confirming that in the “empty” ab3A neuron mutant fly line only the B neuron, which has smaller amplitude spikes (see also Fig. 2A), is present. As expected, the B neuron responds to stimulation with 2-heptanone 1:10,000 vol/vol (mediated by its endogenous *Or85b*, top), similar to wild-type flies (right). **(B)** In the ab3A “empty” neuron mutant, the lack of *Or22a* expression in the A neuron was verified by the lack of spiking upon stimulation with ethyl hexanoate 1:100 vol/vol (left). In wild-type flies (Canton-S), as expected, the A neuron (large spikes) respond strongly to stimulation with ethyl hexanoate. The bottom part of the panel shows the population responses of mutant (left) and wild-type flies (right). Data represents the net number of spikes in response to stimulation (n= 6-7, obtained from 3 females, dots denote individual data points; boxes represent the 25 and 75% quartiles, whiskers represent the 10 and 90% quartiles, and the vertical line inside the boxes indicate the median). Differences between the responses of flies stimulated with the odorant and the control solvent were statistically different in the case of wild-type flies (right, * p<0.05, Wilcoxon-matched pairs test) but not in the case of flies lacking Or22a (left, p>0.05). **(C)** Similarly, lack of Or67d expression in the at1 neuron in the *Or67d*^GAL4^ fly line (“empty at1 neuron” mutant) was verified by the lack of responses to 11-*cis* vaccenyl acetate 1:100 vol/vol (left; ethanol was used as the solvent control). The top shows representative electrophysiological recordings and the bottom the population response (data obtained and represented as explained in A). In wild-type flies, at1 neurons respond strongly to stimulation with this compound (right, * p<0.05).

**Figure supplement 4.**
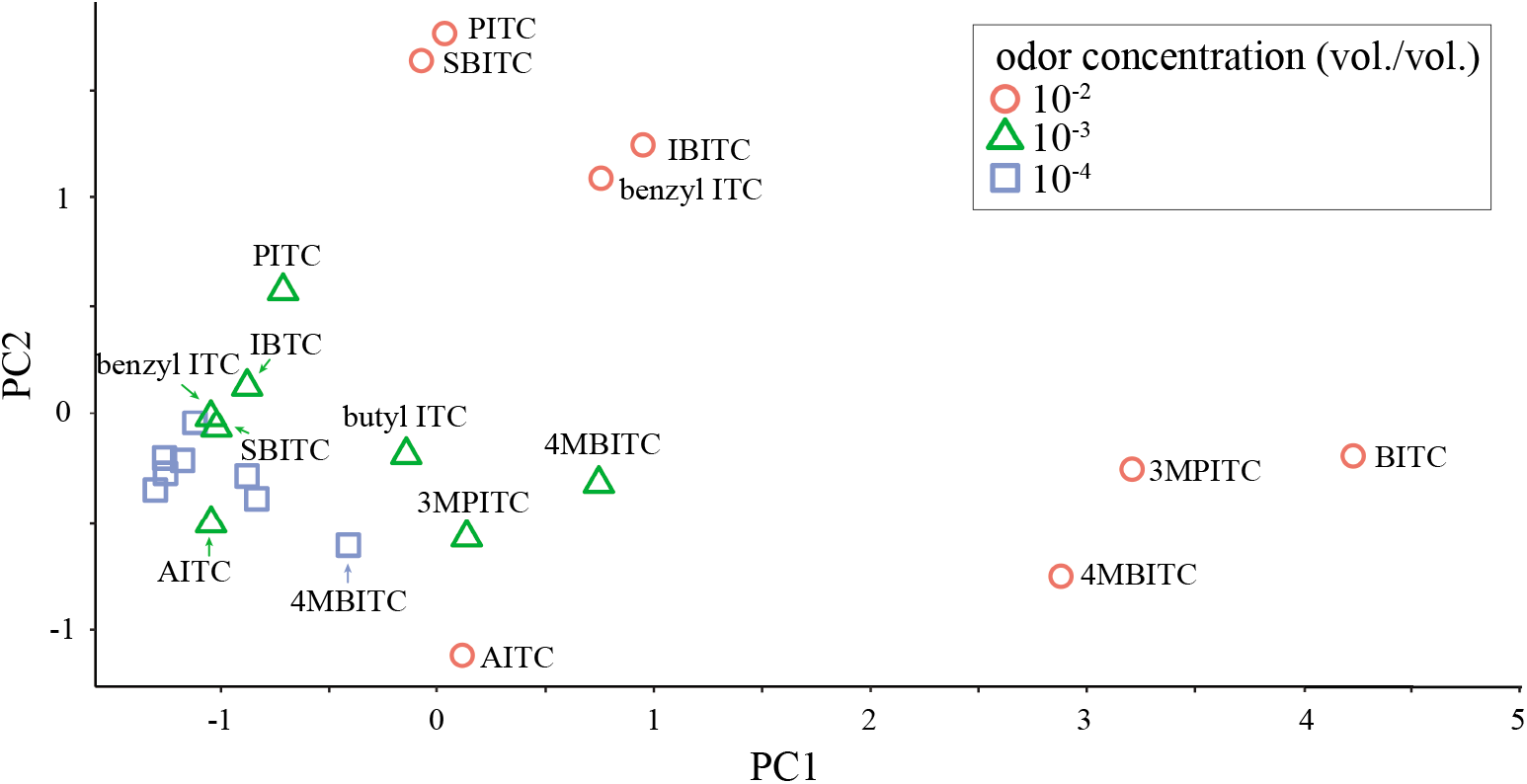
Principal component analysis (PCA) of median responses from the three *S. flava* paralogs. Responses of Or67b paralogs in odor space, generated by PCA of median net spiking responses (control-subtracted) from the three *S. flava* paralogs to eight ITCs tested at 1:100 (red circles), 1:1,000 (green triangles), and 1:10,000 vol/vol (blue rectangles). The axes display the two first principal components (PC1 and PC2) which respectively explain 75% and 17% of the variance.

**Figure supplement 5.**
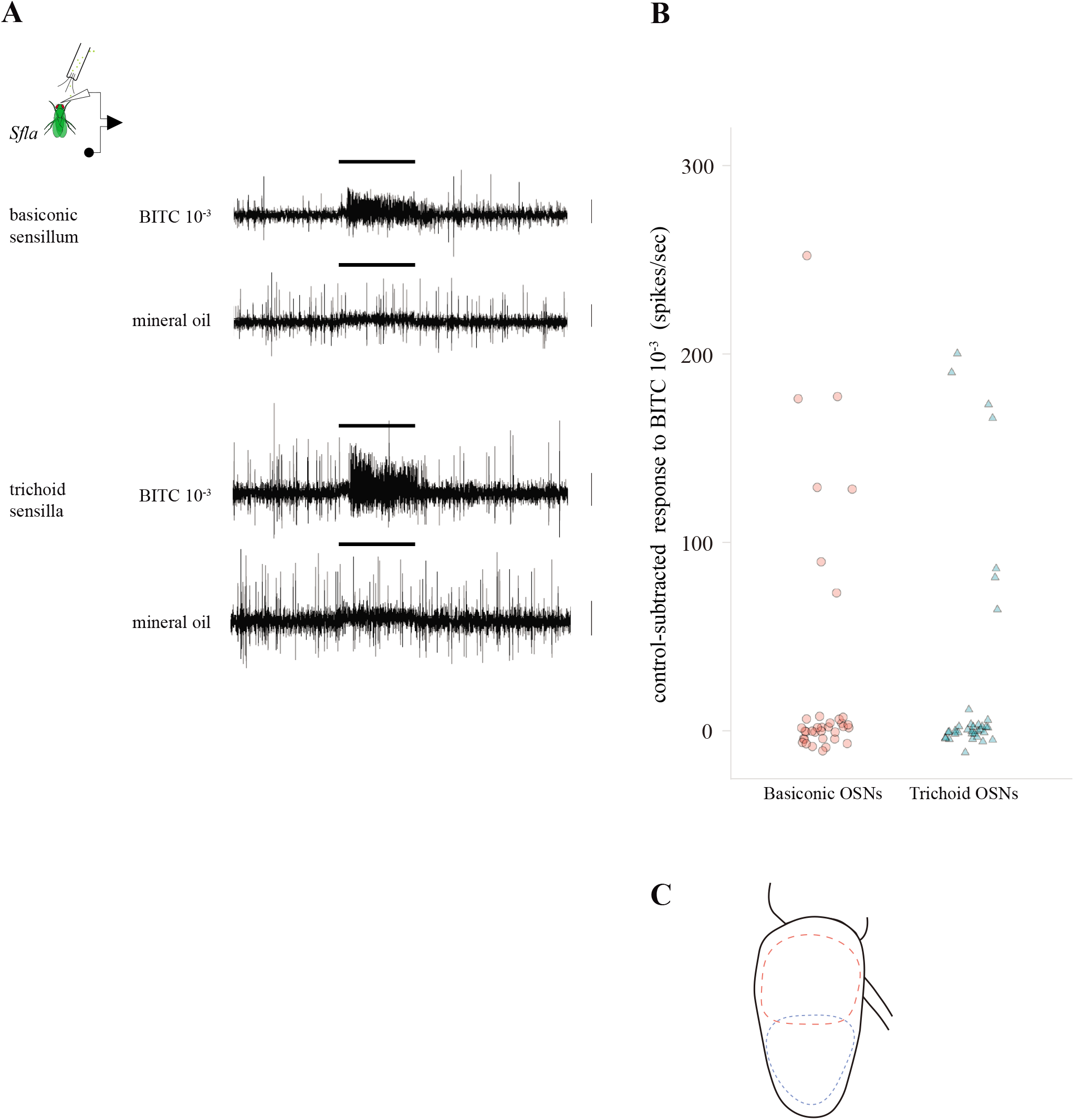
Antennal OSNs respond to ITCs in *S. flava*. **(A)** Representative electrophysiological recordings obtained from antennal *S. flava* OSN housed in basiconic-like (top) and trichoid-like (bottom) sensilla in response to stimulation with BITC 1:1,000 vol/vol and the mineral oil solvent control. In both cases, the OSNs with smaller spike amplitude (“B” neurons) were strongly activated by BITC. **(B)** Responses of *S. flava* individual antennal basiconic-like and trichoid-like OSNs (net number of spikes, control-subtracted; n=36; 3 individuals in each case) in response to BITC 1:1,000 vol/vol. Most basiconic and trichoid OSNs (81%) showed little or no response (median= −10 to 12 net spikes /second), but 19% showed strong responses (range: 74 to 252 and 65 to 200 spikes/second for basiconic and trichoid sensilla, respectively). These results indicate that at least some OSNs are responsive to ITCs in the antenna of *S. flava*, in agreement with the finding that at least the paralogs of Or67b are tuned to these compounds in this fly species. **(C)** Schematic distribution of ITC sensitive OSNs in basiconic-like and trichoid-like sensilla on the antennae. These sensilla distributed proximally and distally (inside of the broken red and blue lines) respectively on the ventral side of antennae.

**Figure supplement 6.**
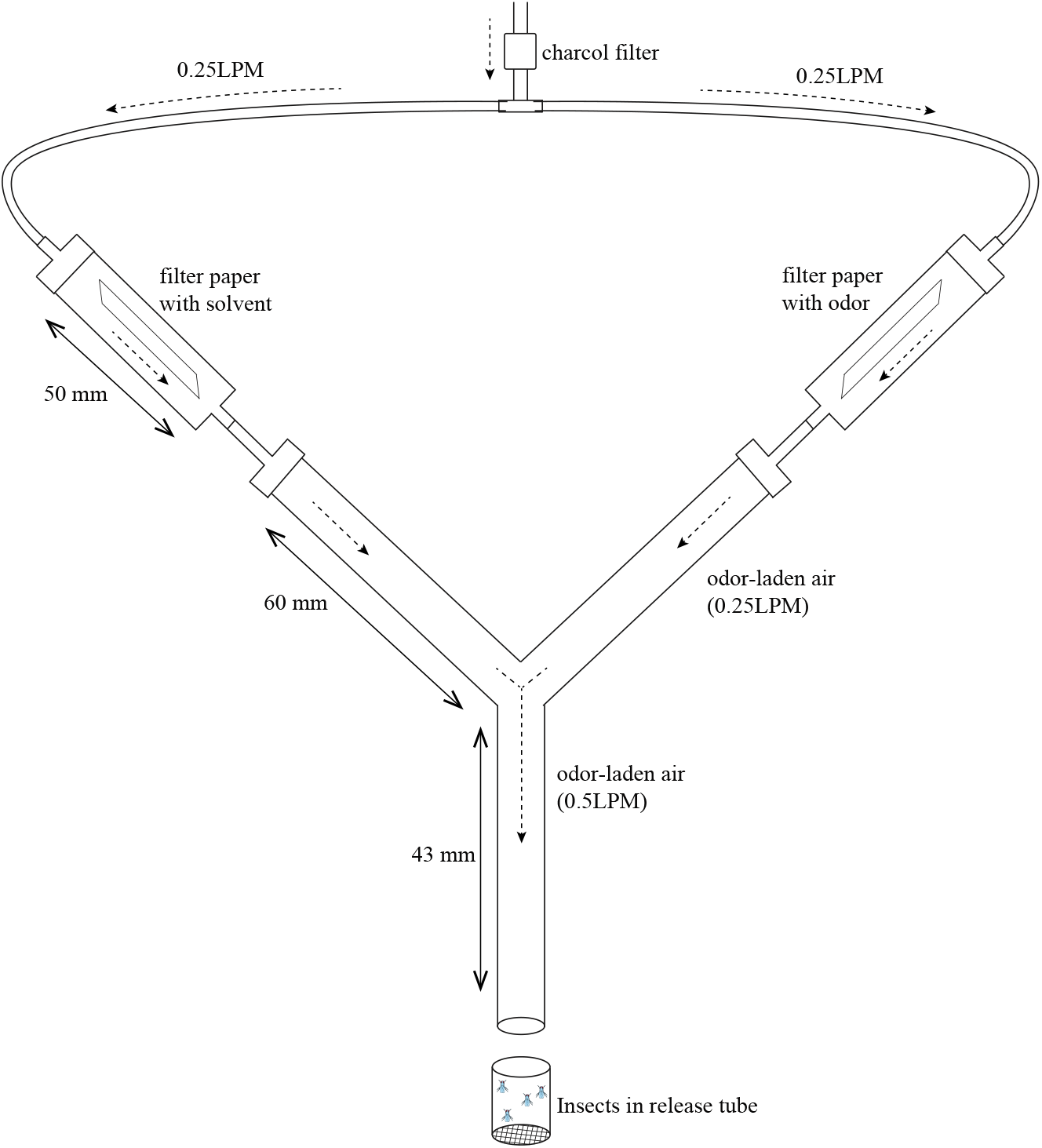
Detailed schematic representation of the device used to test olfactory behavioral responses. The responses of insects were tested using a dual-choice “Y-shaped” olfactometer modified from one previously published *e.g.* ^64^. The “Y” part of the maze was a propylene connector; the open ends of the arms of the connector were each connected to a 1 or a 5 ml plastic syringe containing the odor/odorant stimulus or the control stimulus. Single odorants (or the solvent control) were loaded in a piece of filter paper which was placed in 1 ml syringes; plant leaves (or wet tissue paper as a control) were placed in 5-ml syringes. Charcoal-filtered air was delivered to the maze and adjusted to 0.5 liters per minute (LPM) using a flowmeter; thus, odor airflow in each arm was 0.25 LPM, and at the base of the maze again 0.5 LPM. Three-four insects were placed in individual open-top releasing containers 2 hours before experiments with a piece of tissue paper soaked in distilled water (in the case of *D. melanogaster*) or ca. 20 hours with a cotton piece soaked in honey water solution (in the case of *S. flava*). For experiments in Figure 6C, transgenic flies were placed in individual releasing containers with a piece of tissue paper embedded in water 24 hours before experiments. Before each test, the release container was placed during 45-60 seconds in ice to slow down insect activity. Each test started when the open-top of the insect container was carefully slid into the open end of the long arm of the “Y”. Thus, upon being released, insects could walk upwind towards the “decision point” (intersection of the short and long arms of the “Y”) and turn towards either the odor-laden or the odorless arm of the maze. A choice was considered as such only if the insect walked past at least 1 cm into the arm, orienting upwind. Although 3-4 insects were released in each test (to increase the possibility that at least one insect made a choice), only the first choice (control arm or odorous arm) was recorded. If a second insect made a choice for the other arm within 5-7 seconds of the first insect, the test was discarded. Each test lasted a maximum of five minutes, and each group of insects was used only once. The position of the control and the test arms was switched every one or two tests to control for positional asymmetries. The whole device was illuminated with white light or green light (in the case of tests with *S. flava*).

**Figure supplement 7.**
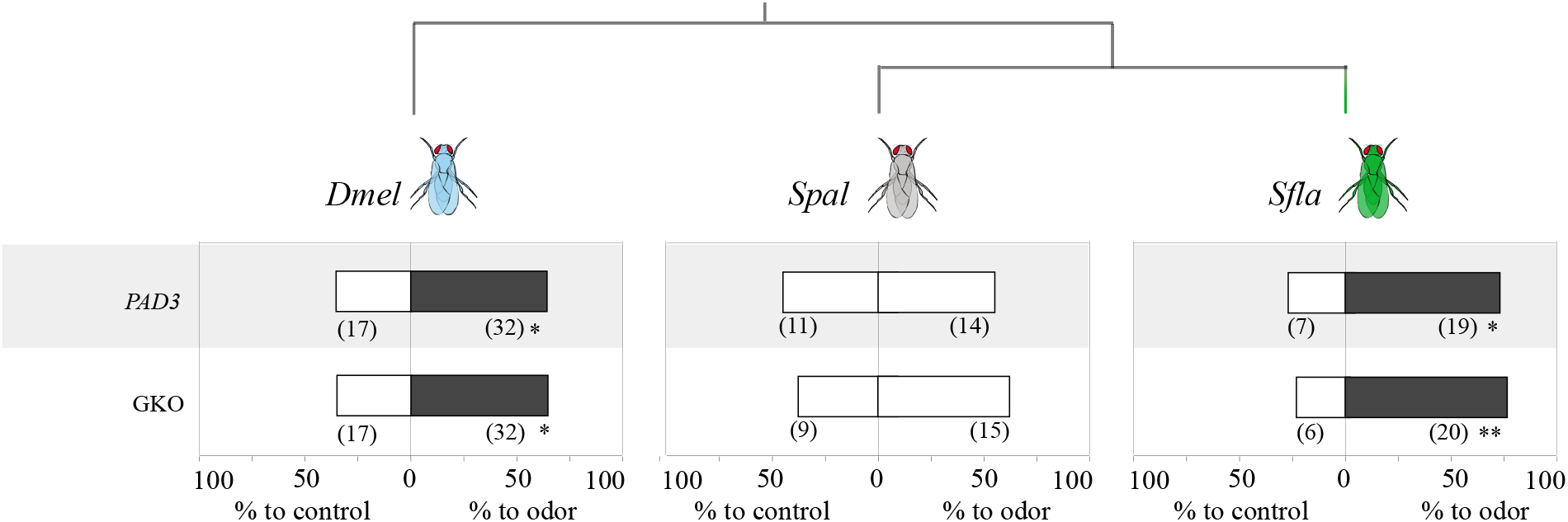
Olfactory behavioral responses of *S. flava* and its microbe-feeding relatives *S. pallida* and *D. melanogaster* to *Arabidopsis*. A dual choice y-maze was used to quantify olfactory behavioral responses of *D. melanogaster*, *S. pallida* and *S. flava* to *PAD3* and GKO *A. thaliana* mutants as described in Figure 5. One arm of the maze offered constant *A. thaliana* odor airflow, while the control arm offered a constant odorless humidified airflow. Data was analyzed using two-tailed Binomial tests (* p<0.05, ** p<0.01). *S. flava* and *D. melanogaster*, but not *S. pallida,* were attracted to leaf VOCs from both *A. thaliana* mutant lines. In this experiment only GKO and *PAD3* plants were used because as explained before, the *PAD3* genotype is a better control for GKO plants than Col-0.

**Figure supplement 8.**
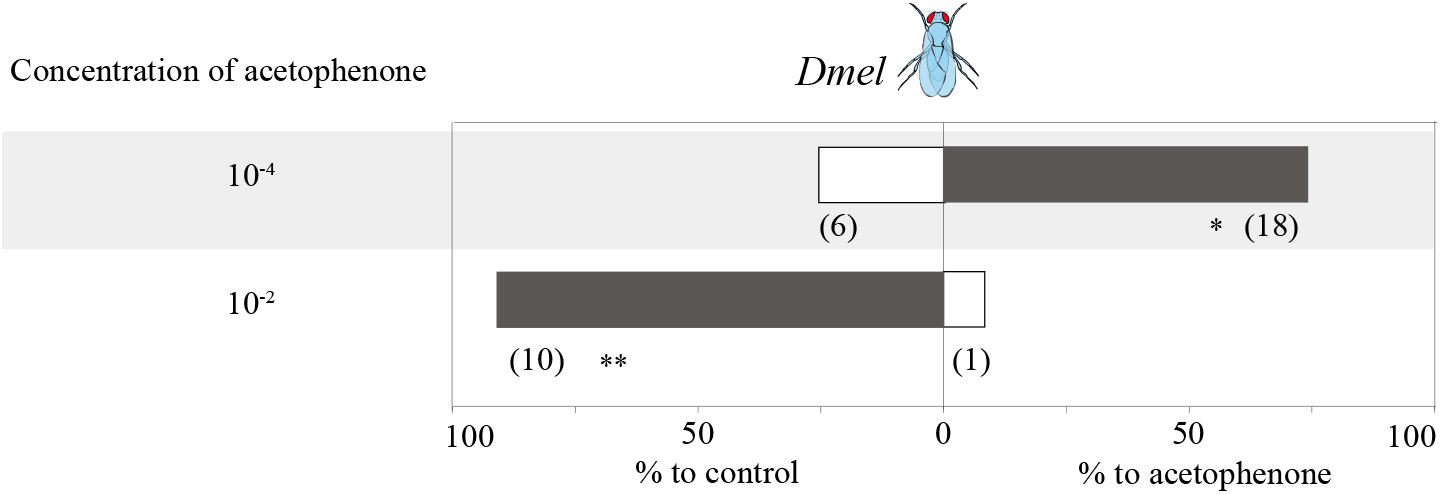
Concentration-dependent behavioral responses of wild-type *D. melanogaster* to acetophenone. Behavioral responses of (strain: Canton-S) flies tested with two concentrations of acetophenone. One arm of the maze offered either acetophenone at 1:10,000 or 1:100 vol/vol, while the other arm had mineral oil. Flies were attracted to the lower concentration but repelled by the higher concentration (Binomial tests, *p<0.05; **p<0.01).

## SOURCE DATA CAPTION

**Source data *Or67bs* are expressed in *Scaptomyza* spp.**

**(A)** Amplification of *Marf* from cDNA generated from whole body extracts of *S. flava* and *S. pallida* adults (+RT). As expected, *Marf* does not amplify in templates treated with DNaseI without reverse transcriptase (– RT), used as a negative control. **(B)** Amplification of *Or67b* genes from *S. pallida* and *S. flava* whole adult cDNA, which reveal *in vivo* transcription of *Or67b* genes in adult *Scaptomyza* (arrow). Ladder (firsts column) is GeneRuler™ 1 kb plus (ThermoFisher, USA).

## SUPPLEMENTARY FILES

**Supplementary file 1 Phylogenetic analysis**

Phylogenetic dataset summary, sequence accession numbers, genome sequence coordinates and phylogenetic model parameters and results.

**Supplementary file 2 Molecular evolution analyses**

Selected parameters and results from branch d*_N_*/d*_S_* tests of all *S. flava* Ors CDS and the expanded *Or67b* dataset.

**Supplementary file 3 Or67b synteny**

The three spreadsheets indicate pGOIs for the three *Or67b* orthologs identified in *S. flava.* For each sheet, we have the following columns: “Position from GOI in *S. flava”* (*e.g.* -1 if one gene upstream of *Or67b,* +2 if two genes downstream of *or67b*); “*S. flava* Annotation ID”; “*D. grimshawi* homolog” (identified via blastn searches); “*D. grimshawi* homolog scaffold”; “*S. pallida* homolog locations”, which include scaffold and coordinates; “*D. melanogaster* homolog”, and “*D. melanogaster* homolog coordinates”. “NA” is written in the cell if homologs were not found after executing blastn, blastx and tblastx searches.

**Supplementary file 4 PCR primers**

Nucleotides in lower case are either in untranslated sequence (CDS amplification) or are restriction enzyme cut sites (RE cut site addition). CDS amplification primers were used to amplify full *Or67b* CDS sequence from cDNA. Primers labeled “RE cut site addition” were used to engineer restriction enzyme cut-sites via PCR mutagenesis in order to ligate *Or67b* CDS sequences into the pUASTattB plasmid. All sequences are listed in a 5’ to 3’ orientation.

**Supplementary file 5 Principal component coordinates**

**Supplementary file 6 pGOI sequences**

**Supplementary file 7 list of chemicals**

